# Engineering the yeast *Saccharomyces cerevisiae* for the production of L-(+)-ergothioneine

**DOI:** 10.1101/667592

**Authors:** Steven A. van der Hoek, Behrooz Darbani, Karolina E. Zugaj, Bala Krishna Prabhala, Mathias Bernfried Biron, Milica Randelovic, Jacqueline B. Medina, Douglas B. Kell, Irina Borodina

**Author notes:** **Correspondence:** Irina Borodina & Douglas B. Kell, The Novo Nordisk Foundation Center for Biosustainability, Technical University of Denmark, Kemitorvet 200, 2800 Kgs. Lyngby, Denmark. Institute of Physics, Chemistry and Pharmacy, Faculty of Science, University of Southern Denmark, Odense, Denmark (Bala Krishna Prabhala).

## Abstract

L-(+)-Ergothioneine is an unusual, naturally occurring antioxidant nutraceutical that has been shown to help reduce cellular oxidative damage. Humans do not biosynthesise it, so can acquire it only from their diet; it exploits a specific transporter (SLC22A4) for its uptake. ERG is considered to be a nutraceutical and possible vitamin that is involved in the maintenance of health, and seems to be at too low a concentration in several diseases *in vivo*. Ergothioneine is thus a potentially useful dietary supplement. Present methods of commercial production rely on extraction from natural sources or on chemical synthesis. Here we describe the engineering of the baker’s yeast *Saccharomyces cerevisiae* to produce ergothioneine by fermentation in defined media. After integrating combinations of ERG biosynthetic pathways from different organisms, we screened yeast strains for their production of ERG. The highest-producing strain was also engineered with known ergothioneine transporters. The effect of amino acid supplementation of the medium was investigated and the nitrogen metabolism of *S. cerevisiae* was altered by knock-out of *TOR*1 or *YIH*1. We also optimized the media composition using fractional factorial methods. Our optimal strategy led to a titer of 598 ± 18 mg/L ergothioneine in fed-batch culture in bioreactors. Because *S. cerevisiae* is a GRAS (‘generally recognised as safe’) organism that is widely used for nutraceutical production, this work provides a promising process for the biosynthetic production of ERG.

## 1 Introduction

Ergothioneine (ERG) (2-mercaptohistidine trimethylbetaine, IUPAC name (2S)-3-(2-Thioxo-2,3-dihydro-1H-imidazol-4-yl)-2-(trimethylammonio)propanoate) is a naturally occurring antioxidant that can be found universally in plants and mammals (Melville, 1959); it possesses a tautomeric structure, but is mainly present in the thione form at physiological pH (figure 1). Ergothioneine was discovered in 1909 in the ergot fungus *Claviceps purpurea* (Tanret, 1909), and its structure was determined two years later (Barger and Ewins, 1911). Subsequently, several other organisms were found to produce ergothioneine, including the filamentous fungus *Neurospora crassa* (Genghof et al., 1956), the yeast *Schizosaccharomyces pombe* (Pluskal et al., 2014), various actinobacteria (Genghof, 1970) including *Mycobacterum smegmatis* (Seebeck, 2010), and in particular various basidiomycetes (mushrooms) (Genghof, 1970).

**Figure 1:**
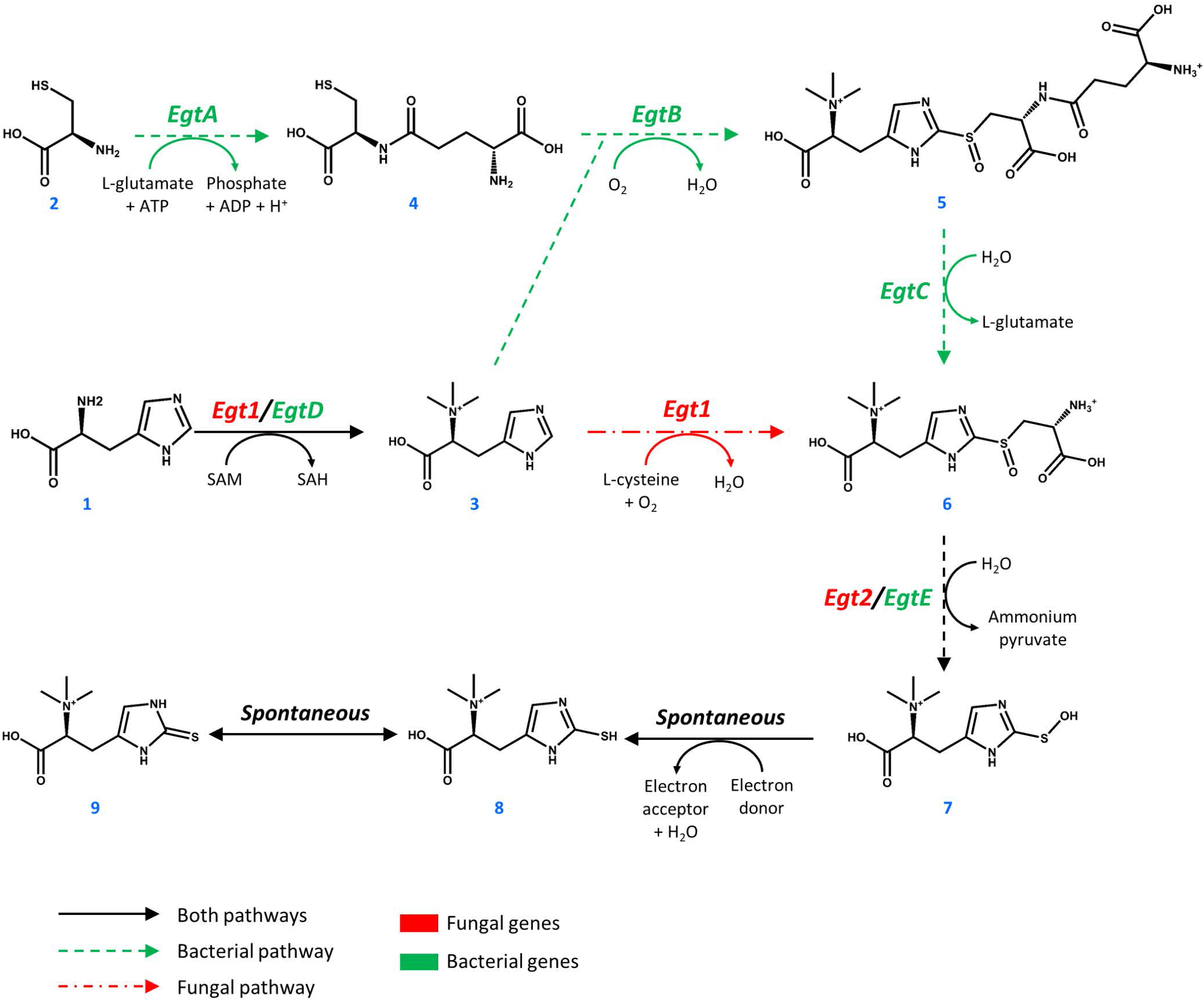
Pathways towards ergothioneine biosynthesis in bacteria and fungi.

Ergothioneine is synthesized from one molecule of L-histidine (**1**), one molecule of L-cysteine (**2**), and 3 methyl groups donated by S-adenosyl-L-methionine (SAM, figure 1). In *M. smegmatis*, the reaction sequence is catalyzed by five enzymes, encoded by *egtA-E* genes arranged in an operon (Sao Emani et al., 2018). Four enzymes of the cluster EgtA, EgtB, EgtC, and EgtD catalyze four individual reactions that methylate L-histidine to form hercynine (**3**), convert L-cysteine to γ-L-glutamyl-L-cysteine (**4**), combine hercynine and γ-L-glutamyl-L-cysteine to generate γ-L-glutamyl-S-(hercyn-2-yl)-L-cysteine S-oxide (**5**) and produce the S-(hercyn-2-yl)-L-cysteine S-oxide (**6**, HCO). In fungi, the biosynthetic pathway is different, as a single enzyme Egt1 catalyzes the methylation of L-histidine (**1**) to give hercynine (**3**), which in turn is sulfoxidized with cysteine, producing HCO (**6**). HCO is converted into 2-(hydroxysulfanyl)hercynine (**7**) by a β-lyase, encoded by *egtE* in *M. smegmatis* and by the *EGT2* gene in fungi. This compound is spontaneously reduced to ergothioneine (**8**, thiol form and **9**, thione form).

The antioxidant properties of ERG include the scavenging of free radicals and reactive oxygen species (Akanmu et al., 1991; Park et al., 2010; Ta et al., 2011) and the chelation of divalent metal ions (Hanlon, 1971). ERG has been shown to reduce oxidative damage in a variety of mammals (Cheah and Halliwell, 2012; Deiana et al., 2004). In humans, ERG is mainly accumulated in the liver, the kidneys, in erythrocytes, the eye lens, and in seminal fluid (Leone and Mann, 1951; Melville et al., 1954; Shires et al., 1997). It is transported by a specific transporter SLC22A4 (previously known as OCTN1) (Grundemann et al., 2005; Tschirka et al., 2018). The natural selection of such a transporter implies that ergothioneine is involved in the maintenance of health or the mitigation of disease, and it may even be a vitamin (Paul and Snyder, 2010). ERG has demonstrated effects in *in vivo* models of several neurodegenerative diseases (Link, 1995; Song et al., 2014; Yang et al., 2012), in ischaemia reperfusion injury (Bedirli et al., 2004; Sakrak et al., 2008a, 2008b), and in a variety of other diseases (Halliwell et al., 2018). Ergothioneine accumulates at sites of injury through the upregulation of SLC22A4/OCTN1 (Cheah et al., 2016; Tang et al., 2016). It is only slowly metabolized and excreted in humans (Cheah et al., 2017).

Because humans cannot produce ERG, they must obtain it through their diet. Although plants and animals also accumulate it to some degree, the main natural dietary source of ERG is basidiomycete mushrooms, where some species contain up to 7 mg of ERG per gram dry weight (Ey et al., 2007; Halliwell et al., 2018; Kalaras et al., 2017; Pfeiffer et al., 2011). Because of the beneficial effects and possible involvement of ERG in disease, ergothioneine may potentially prove its value in the global dietary supplement market, which was estimated at some $241.1 billion in 2019 (Wang et al., 2016). Currently, commercial ergothioneine is extracted from mushrooms or synthesized chemically. Production of ergothioneine in microbial cell factories would provide a sustainable low-cost alternative to its current manufacturing processes. So far, fermentative ERG production has been reported in bacteria and filamentous fungi, including *Methylobacterium aquaticum* strain 22A (Alamgir et al., 2015), *Aureobasidium pullulans* (Fujitani et al., 2018), *Rhodotorula mucilaginosa* (Fujitani et al., 2018)*, cyanobacteria* (Pfeiffer et al., 2011), *Aspergillus oryzae* (Takusagawa et al., 2019) and *Escherichia coli* (Osawa et al., 2018; Tanaka et al., 2019). To the best of our knowledge, there are no reports on ergothioneine production in baker’s yeast, which is the preferred host for the production of nutraceuticals (Huang et al., 2008; Li and Borodina, 2014; Yuan and Alper, 2019). In this study we describe the metabolic engineering of the yeast *S. cerevisiae* for the production of ergothioneine, reaching a titer of 0.6 g/L in fed-batch fermentation.

## 2 Materials and Methods

### 2.1 Strains and chemicals

*S. cerevisiae* strain CEN.PK113-7D (MATa URA3 HIS3 LEU2 TRP1 MAL2-8^c^ SUC2) was a gift from Peter Kötter (Goethe University, Frankfurt/Main, Germany). *Schizosaccharomyces pombe* strain DSM 70572, obtained from Leibniz-Institut DSMZ-Deutsche Sammlung von Mikroorganismen und Zeilkulturen GmbH (Germany) was used for genomic DNA extraction. *E. coli* DH5α was used for cloning. Ergothioneine (catalogue #E7521, ≥98% purity) was from Sigma-Aldrich, hercynine (catalogue # H288900, ≥95% purity) was from Toronto Research Chemicals Inc. Simulated fed-batch medium components (EnPump 200) were from EnPresso GmbH (Germany). These components consist of a polysaccharide, in powder form, to add to the medium and an enzyme to release glucose from the polysaccharide. The rate of release is dependent on the enzyme concentration and allows for simulated carbon-limited fermentation.

### 2.2 Cloning

CpEgt1, MsEgtA and MsEgtE were codon-optimized for *S. cerevisiae* using GeneGenie (Swainston et al., 2014), while the other genes were codon-optimized for *S. cerevisiae* using the codon-optimization tool provided by GeneArt. The genes were then ordered as synthetic gene strings from IDT DNA (MsEgtA) or GeneArt (all other synthetic gene strings). The only exceptions were two genes from *Schizosaccharomyces pombe*, which were isolated from genomic DNA. The DNA sequences of the genes are in supplementary table 1. All yeast strain construction was done using CRISPR/Cas9 and EasyClone-MarkerFree methods (Jessop-Fabre et al., 2016; Stovicek et al., 2015). Correct cloning was validated by sequencing (Eurofins Genomics). Correct genome modification of yeasts was validated by colony PCR. The details on primers (supplementary table 2 and 3), biobricks (supplementary table 4), plasmids (supplementary table 5 and 6), and strains (supplementary table 7) are in supplementary materials.

### 2.3 Media and small-scale cultivation conditions

For *E. coli* selection, we used Lysogeny Broth (LB) with 100 mg/L ampicillin. For the selection of yeast strains, we used Yeast-Peptone-Dextrose (YPD) agar supplemented with 200 mg/L G418 for selection of Cas9 vector and 100 mg/L nourseothricin for selection of the gRNA vector. Synthetic Complete (SC) medium was prepared using 6.7 g/L Yeast Nitrogen Base without amino acids from Sigma-Aldrich, 1.92 g/L Synthetic Drop-out supplement without histidine from Sigma-Aldrich and 76 mg/L histidine. For ERG production, yeast strains were cultivated in SC medium with 20 or 40 g/L glucose or with 60 g/L EnPump substrate (and 0.6% enzyme reagent) as carbon source. Precultures for cultivation experiments were prepared by inoculating a single colony of a strain into 5 ml of the medium used in the cultivation experiment and incubating at 30 °C and 250 rpm for 24 hours. The cultivations were performed in 24-deep-well plates from EnzyScreen, 3 ml of medium was used per well and the starting OD600 was 0.5. The plates were incubated at 30 °C with 250 rpm agitation.

### 2.4 HPLC analyses

To determine the extracellular ergothioneine concentration, a 1-ml sample of fermentation broth was centrifuged at 3,000 x g for 5 min, the supernatant was moved into an HPLC vial and stored at −20 °C until the analysis. The remaining cell pellet was washed twice with 1 ml MilliQ water and resuspended in 1 ml water. The extraction of intracellular ERG was performed according to Alamgir et al. (Alamgir et al., 2015) as following. The mixture was heated at 94° C for 10 minutes, vortexed at 1,600 rpm for 30 minutes using a DVX-2500 Multi-Tube Vortexer from VWR, and centrifuged at 10,000 × g for 5 minutes. The supernatant was transferred into an HPLC vial and stored at +4 °C until analysis. The ERG concentrations were measured using a Dionex Ultimate 3000 HPLC system. Quantification was done based on standard curves using Chromeleon software. 5 ◻l of sample was injected on a Cortects UPLC T3 reversed-phase column (particle size 1.6 μm, pore size 120 Å, 2.1 × 150 mm). The flow rate was 0.3 ml/min, starting with 2.5 minutes of 0.1% formic acid, going up to 70% acetonitrile, 30% 0.1% formic acid at 3 minutes for 0.5 minutes, after which 100% 0.1% formic acid was run from minute 4 to 9. Ergothioneine was detected at a wavelength of 254 nm. For analysis of bioreactor samples, we additionally quantified glucose, ethanol, pyruvate, and acetate concentrations by HPLC as described (Borodina et al., 2015).

### 2.5 Fed-batch fermentation in bioreactors

A single colony from a YPD plate with ST8927 colonies (see below) was used to inoculate 5 ml of minimal media in 13-ml tube. The tube was incubated at 30° C and 250 rpm overnight. This overnight culture was transferred into 95 ml mineral medium in 500 ml baffled shake flask. The shake flask was then incubated overnight at 30° C and 250 rpm. 40 ml of this dense culture was used to inoculate 60 ml mineral medium in a new 500 ml baffled shake flask. Two shake flasks were prepared this way. These shake flasks were incubated at 30° C and 250 rpm for 4 hours, the content of both shake flasks was combined, then centrifuged at 3,000 x g for 5 min. The supernatant was discarded, the pellet was washed with 25 ml sterile water, resuspended and centrifuged as before. The supernatant was discarded and the pellet resuspended in 10 ml mineral medium. This was then used to inoculate 0.5 l mineral medium in a 1 l Sartorius bioreactor. The starting OD600 was 0.85. The stirring rate was set at 500 rpm, the temperature was kept at 30° C, and pH was maintained at pH 5.0 using 2 M KOH and 2 M H_2_SO_4_. The feeding was started as soon as CO_2_ in the off-gas decreased by 50%. The initial feed rate was set at 0.6 g glucose h^−1^, linearly increasing to 2.5 g glucose h^−1^ over the span of 25.5 hours. After that, the feed was set at a constant 1.4 g glucose h^−1^ and 17.8 hours later, the feeding rate was set to a constant 2.9 g glucose h^−1^. The feed was stopped at 84 hours. At 60.5 and 75.5 hours, 2 g (NH_4_)_2_SO_4_ was added as a sterile 100 g/l solution. At 60.5 and 73.5 hours, 0.5 g MgSO_4_ was added as a sterile 50 g/l solution, while 4 ml sterile trace metals solution and 2 ml sterile vitamin solution were added.

### 2.6 Propidium iodide staining and flow cytometry analysis

Precultures were prepared by inoculating a single colony of strain ST7574, ST8461 and ST8654 into separate 14-ml tubes containing 5 ml of SC + 40 g/L glucose + 1 g/L His/Cys/Met and incubating at 30 °C and 250 rpm for 24 hours. Precultures were used to inoculate 25 ml SC + 40 g/L glucose + 1 g/L His/Cys/Met at a starting OD_600_ of 0.5, which was incubated at 30 °C and 250 rpm for 72 hours. Every 24 hours, a 1 ml sample of cell culture was taken from the yeast cultivation. This sample was washed two times with 1 ml phosphate-buffered saline (PBS), subsequently resuspended in 0.5 μg/ml propidium iodide (PI) in PBS and incubated for 20 minutes at room temperature in the dark. After incubation, the cells were washed two times with PBS and the percentage of PI stained cells was determined using a MACSQuant VYB system (Miltenyi Biotec). Data analysis was performed using the FlowJo software.

### 2.7 Fluorescent microscopy

Precultures were prepared by inoculating a single colony into a 14-ml tube containing 5 ml YPD medium and incubating at 30 °C and 250 rpm overnight. Overnight precultures of strains containing transporters linked to GFP and the control strain were used to inoculate 5 ml YPD medium in 14 ml tubes at OD_600_ = 2 and were cultured at 30° C and 250 rpm for 5 hours. 1 ml of the culture was harvested at 3,000 × g for 5 minutes. The cells were washed two times with PBS and subsequently pictures of the cells were taken using a Leica DM 400 B system, using Leica Application Suite V4 as image software.

### 2.8 Medium optimization

A two-level fractional factorial based on the components of SC medium, with the levels high (+1, component 5 folds higher than original SC medium) and low (−1, component 5 folds lower than original SC medium), was used to determine the impact of individual components on the yield of ergothioneine. Two different stocks of all the individual components were prepared, one each for high and low concentrations and these were mixed together for all components to yield 64 different designed media. The design matrix for the fractional factorial grid has been attached (supplementary table 8). Precultures were prepared by inoculating a single colony of strain ST8461 into a 14-ml tube containing 5 ml SC medium and incubating at 30 °C and 250 rpm overnight. Precultures were then used to inoculate 300 μl media in 96-deep-well plates from EnzyScreen at OD_600_ = 0.1 in duplicate for each different medium and incubated at 30° C and 225 rpm for 48 hours. Samples for analysis were taken at 24 and 48 hours. Amounts of total ergothioneine were analyzed my LC-MS in MRM mode. The analysis was performed on a Bruker EVOQ (QqQ) coupled to UPLC. The LC part of the LC-MS/MS system consisted of a CTC autosampler module, a high pressure mixing pump and a column module (Advance, Bruker, Fremont, CA, USA). The injection volume was 1 μl. The chromatography was performed on a ZIC-cHILIC column, 150mm × 2.1mm, 3μm particle size, (SeQuant, Merck Millipore), equipped with a 0.5 μ depth filter (KrudKatcher Classic, Phenomenex). Eluent A was 20 mM ammonium acetate, pH adjusted to 3.5 with formic acid in MilliQ water. Eluent B was acetonitrile. The total flow rate of eluent A and B was 0.4 ml/min. The isocratic elution was 30% A, and the total run time was 5min. Retention time was 1.8 min for Ergothioneine. The mass spectrometer was operated with electrospray in the positive ion mode (ESI+). The spray voltage was set to 4500 V. The cone gas flow was 20 l/h, and the cone temperature was set at 350 °C. The heated probe gas flow was set at 50l/h with a temperature of 350 °C. Nebulizer flow was set at 50l/h, and the exhaust gas was turned on. Argon was used as collision gas at a pressure of 1.5 mTorr. Detection was performed in multiple reacting monitoring (MRM) mode. The quantitative transition was 198→95 for Ergothioneine and the qualitative transition was 198→154. The collision energy was optimized to 15 and 7eV respectively. The xms files were converted into cdf files and were analyzed using Mzmine 2.33.

## 3 Results

### 3.1 Expression of bacterial, fungal, and chimeric biosynthetic pathways towards ergothioneine in S. cerevisiae

The biosynthetic genes for ERG production were of both bacterial origin (*M. smegmatis*) and of fungal origin (*Claviceps purpurea*, *Neurospora crassa*, *S. pombe)*. The Egt1 homologues in *C. purpurea* and *S. pombe* were identified by BLASTp using the sequence of Egt1 for *N. crassa* (Genbank accession: XP_956324.3). Similarly, Egt2 from *S. pombe* (Genbank accession: NP_595091.1) was used to find the Egt2 homologues in *N. crassa* and *C. purpurea*. Genbank accession numbers are provided in supplementary table 1. The genes were combined into sixteen pathway variants, where nine pathway variants were made of fungal genes, one pathway variant comprised bacterial genes only, and six variants contained both fungal and bacterial genes. The sixteen yeast strains with different pathway variants were cultivated in three different media and the intra- and extracellular concentrations of ergothioneine were measured (figure 2).

**Figure 2:**
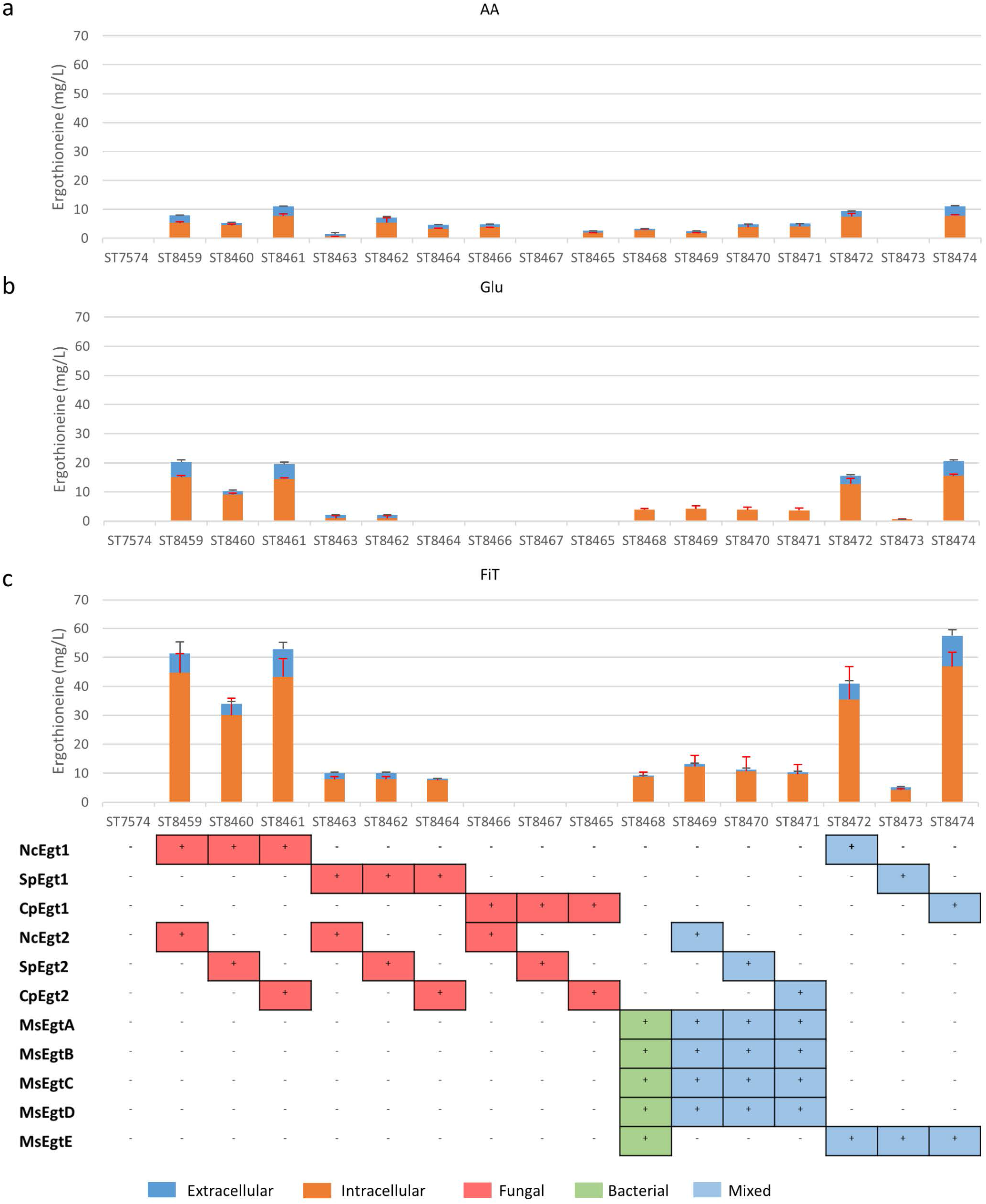
Ergothioneine production in strains with different ERG pathway combinations on three variations of synthetic complete medium: **(A)** AA: 20 g/l glucose and 1 g/l of each L-His, L-Met, and L-Cys, **(B)** Glu: 40 g/l glucose, **(C)** FiT: 60 g/l EnPump substrate for slow glucose release to simulate fed-batch conditions.

ERG titers were the highest in simulated fed-batch medium (up to 60 mg/L as compared to a maximum of 20 mg/L under batch conditions). Of the five best-performing pathway variants, four contained Egt1 from *Neurospora crassa* and any other of the four enzymes catalyzing the 2^nd^ enzymatic step (MsEgtE, NcEgt2, SpEgt2, or CpEgt2). The fifth contained CpEgt1 and MsEgtE. Curiously, CpEgt1 combined with any fungal Egt2 variants only produced ERG on the medium that contained high levels of histidine, cysteine and methionine (figure 2, a). Overall, only 5-30% of the total ERG was secreted, while the rest was retained intracellularly. The strain ST8461, combining Egt1 from *N. crassa* and Egt2 from *C. purpurea,* was selected for further engineering.

### 3.2 Engineering ergothioneine transport

The concentration of ERG inside the cells can be estimated to be ~3.5 mM (for strain ST8461 on simulated fed-batch medium), which is 80-fold higher than in the broth. High intracellular concentrations of a product may inhibit its own biosynthesis or lead to product degradation (Borodina, 2019; Kell, 2019; Kell et al., 2015). We speculated that expressing an ERG-specific transporter might improve the secretion of ERG from the yeast cells. As *M. smegmatis* is known to secrete ergothioneine to levels up to 4 times the intracellular concentration (Sao Emani et al., 2013), we speculated that it may contain an equilibrative or effluxing ERG transporter. On the *M. smegmatis* genome, adjacent to the *egtA-E* operon, there is an open reading frame MSMEI_6084, encoding a protein annotated as a chloramphenicol exporter (supplementary figure 1). The protein has 12 transmembrane domains as predicted by Phyre2 (Kelley et al., 2015) (supplementary figure 2). We expressed codon-optimized variants of this gene and of the (normally concentrative) human ERG transporter SLC22A4 in strain ST8461 (figure 3). However, there was no significant increase of intracellular or extracellular ERG production. To determine why neither transporter had an effect, we investigated their cellular localization by green fluorescent protein (GFP) tagging on the C- or N-termini (figure 3). Interestingly, MsMEI_6084 mainly localized to the vacuolar membrane of *S. cerevisiae*, while human SLC22A4X was only weakly expressed and its localization could not be determined. Since neither of the transporters localized to the plasma membrane specifically, it can explain the lack of effect on ERG secretion.

**Figure 3:**
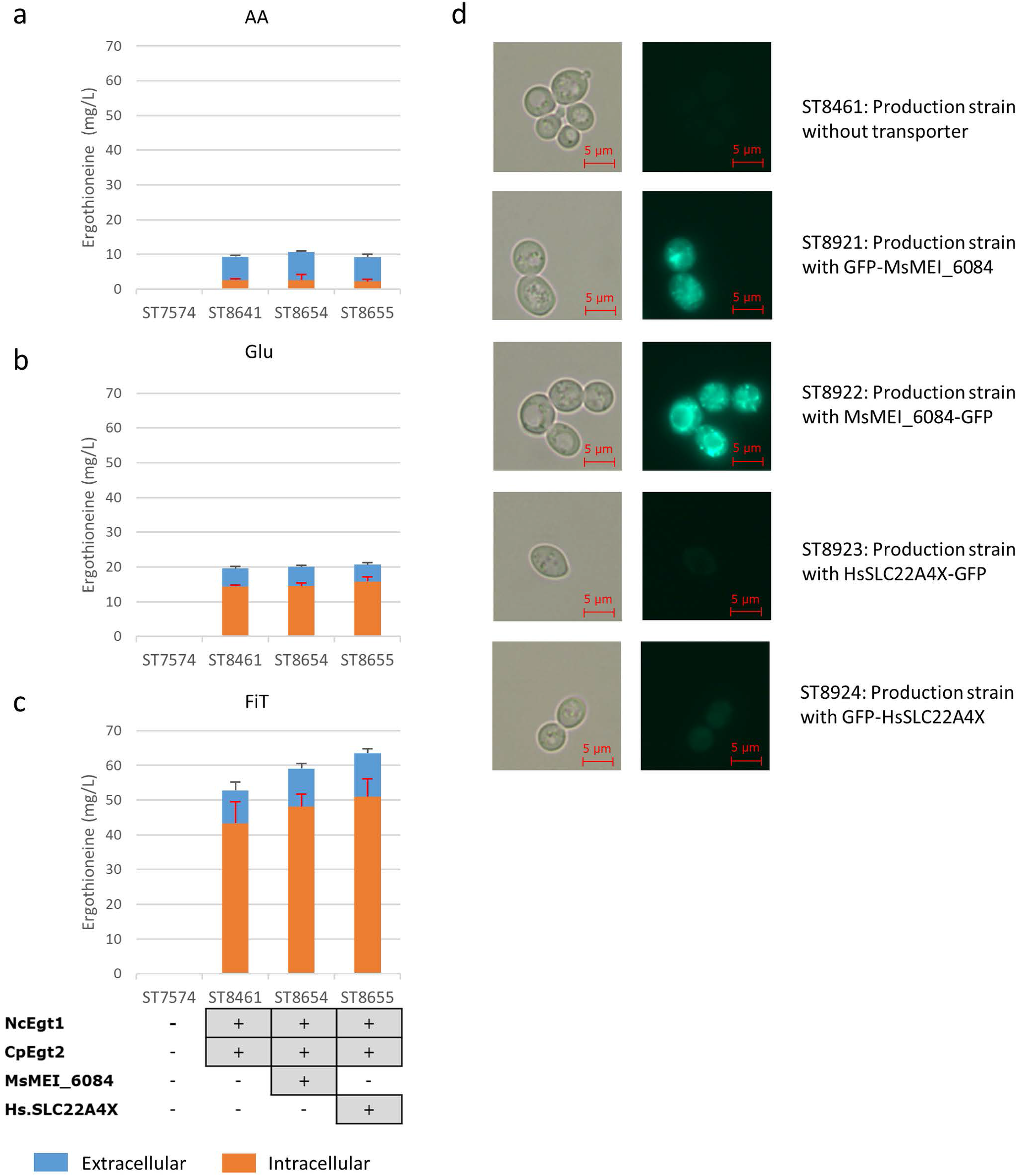
Ergothioneine production by yeast strains expressing human ERG transporter SLC22A4X or a putative ERG transporter from *M. smegmatis* on three variations of synthetic complete medium: **(A)** AA: 20 g/l glucose and 1 g/l of each L-His, L-Met, and L-Cys, **(B)** Glu: 40 g/l glucose, **(C)** FiT: 60 g/l EnPump substrate for slow glucose release to simulate fed-batch conditions, **(D)** Microscope images of yeast cells with ergothioneine transporters linked to GFP. To the left are Brightfield images and to the right GFP images. Top panels; ST8461, ergothioneine producing strain without transporter. Middle-top panels, ST8921, ergothioneine producing strain with a putative transporter from *Mycobacterium smegmatis* linked to GFP at the N-terminus. Middle panels, ST8922, ergothioneine producing strain with a putative transporter from *Mycobacterium smegmatis* linked to GFP at the C-terminus. Middle-bottom panels, ST8923, ergothioneine producing strain with a transporter from *Homo sapiens* linked to GFP at the C-terminus. Bottom panels, ST8924, ergothioneine producing strain with a transporter from *Homo sapiens* linked to GFP at the N-terminus.

### 3.3 Engineering of nitrogen metabolism

The amino acid metabolism of *Saccharomyces cerevisiae* is tightly regulated and its networks highly intertwined (Hinnebusch, 1988, 2005; Hinnebusch and Natarajan, 2002; Ljungdahl and Daignan-Fornier, 2012). Therefore, it would be easier to increase the total amino acid pool rather than increasing the individual amino acid pools. The general amino acid control of yeast is mainly regulated by *GCN4* (Hinnebusch, 1988; Ljungdahl and Daignan-Fornier, 2012) and upregulation of *GCN4* leads to transcription of the biosynthetic genes of various amino acids. As both Tor1 (Hinnebusch, 2005; Ljungdahl and Daignan-Fornier, 2012) and Yih1 (Hinnebusch, 2005) inhibit Gcn2p, a positive regulator of *GCN4*, deletion of either of these enzymes could lead to increased ERG production. In figure 4, we show the results of these alterations in the nitrogen metabolism of *S. cerevisiae* in different media. Yih1 deletion does not have an effect on the production of ergothioneine, while Tor1 only seems to lead to an increase in ERG under batch conditions. Since eventual ERG production is likely to be carried out under fed-batch conditions and neither of the deletions gave a positive effect under fed-batch conditions, we decided not to proceed with these genetic modifications.

**Figure 4:**
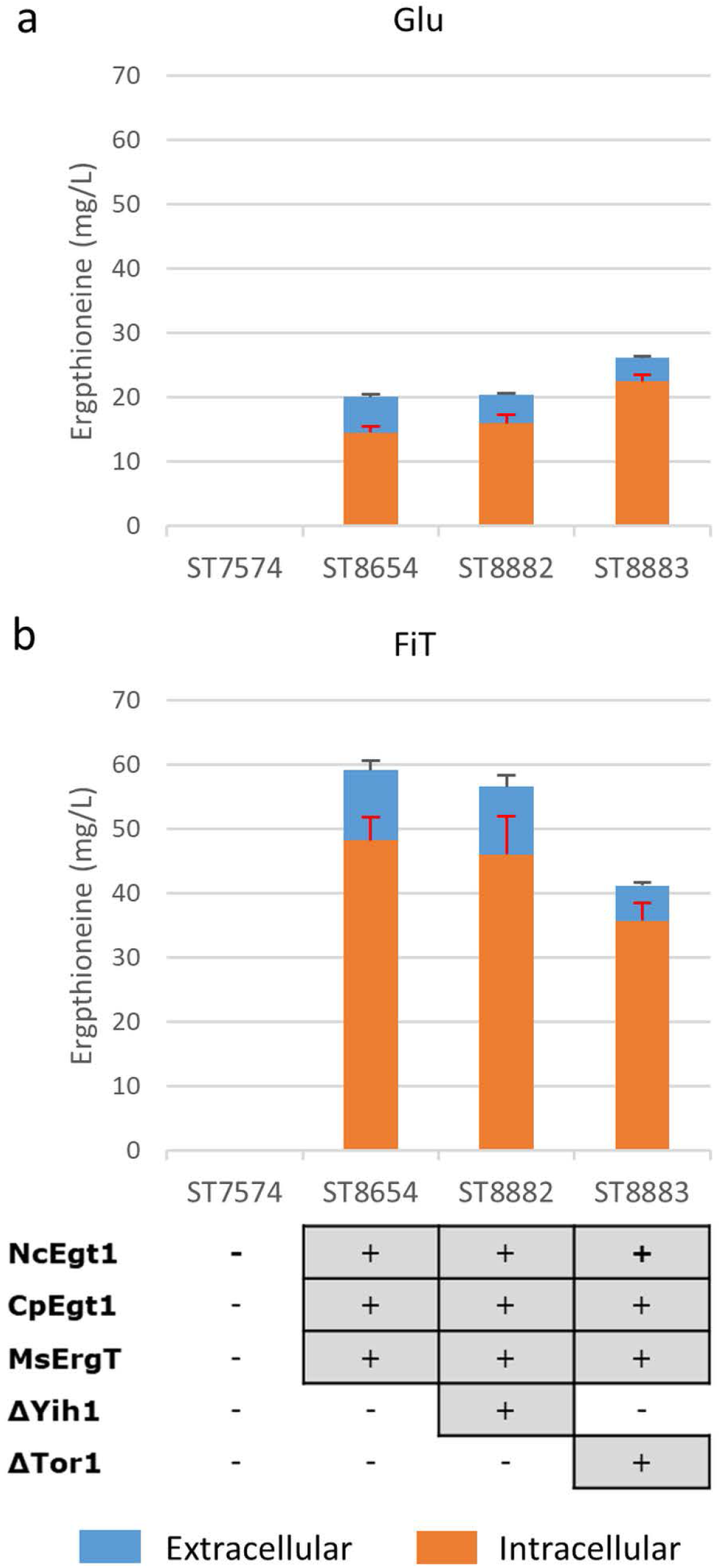
The effect of gene knock-outs linked to nitrogen metabolism on the production of ergothioneine in the production strain with the MsErgT transporter in different media. Genomic alterations are shown as well. **(A)** Glu: SC + 40 g/l glucose **(B)** FiT: SC + 60 g/l EnPump substrate, 0.6% reagent A.

### 3.4 Supplementation of the medium with precursor amino acids

To determine whether the supply of the three amino acids (L-histidine, L-cysteine, and L-methionine) that serve as precursors for ERG biosynthesis is limited, we cultivated several yeast strains with supplementation of 1 g/L or 2 g/L of each of L-methionine, L-cysteine and L-histidine. We chose two producing strains, ST8461 and ST8654, the latter containing the MsMEI_8064 gene. A non-producing strain ST7574 was used as a control. The experiments were performed in shake flasks and growth and ERG production were monitored over the course of three days (figure 5). ERG accumulated primarily in the first 24 hours of cultivation, which would correspond to the exponential growth on glucose, reaching ca. 16 mg/L in both producing strains, independent of any amino acid supplementation. The supplementation, however, affected the cellular growth, with the final OD being approximately 46% and 52% lower when 1 g/L or 2 g/L respectively of the amino acids were added. No degradation of ERG was observed; however surprisingly, there was a large variation in the intracellular vs extracellular distribution of ERG depending on the addition of amino acids. Specifically, the addition of amino acids promoted the excretion of ERG in the stationary phase. We hypothesized that this was due to cell death because of the toxic effects of the added amino acids, in particular histidine (Watanabe et al., 2014) and cysteine (Kumar et al., 2006). Indeed propidium iodide staining of cells sampled every 24 hours for 72 hours, showed an increase in the fraction of dead cells from 9 to 70%, when amino acids were added at concentrations of 1 g/L (supplementary figure 3). Clearly, the concentrations of amino acids used were too high and hence we decided to undertake a more systematic medium optimization approach as described in the next section.

**Figure 5:**
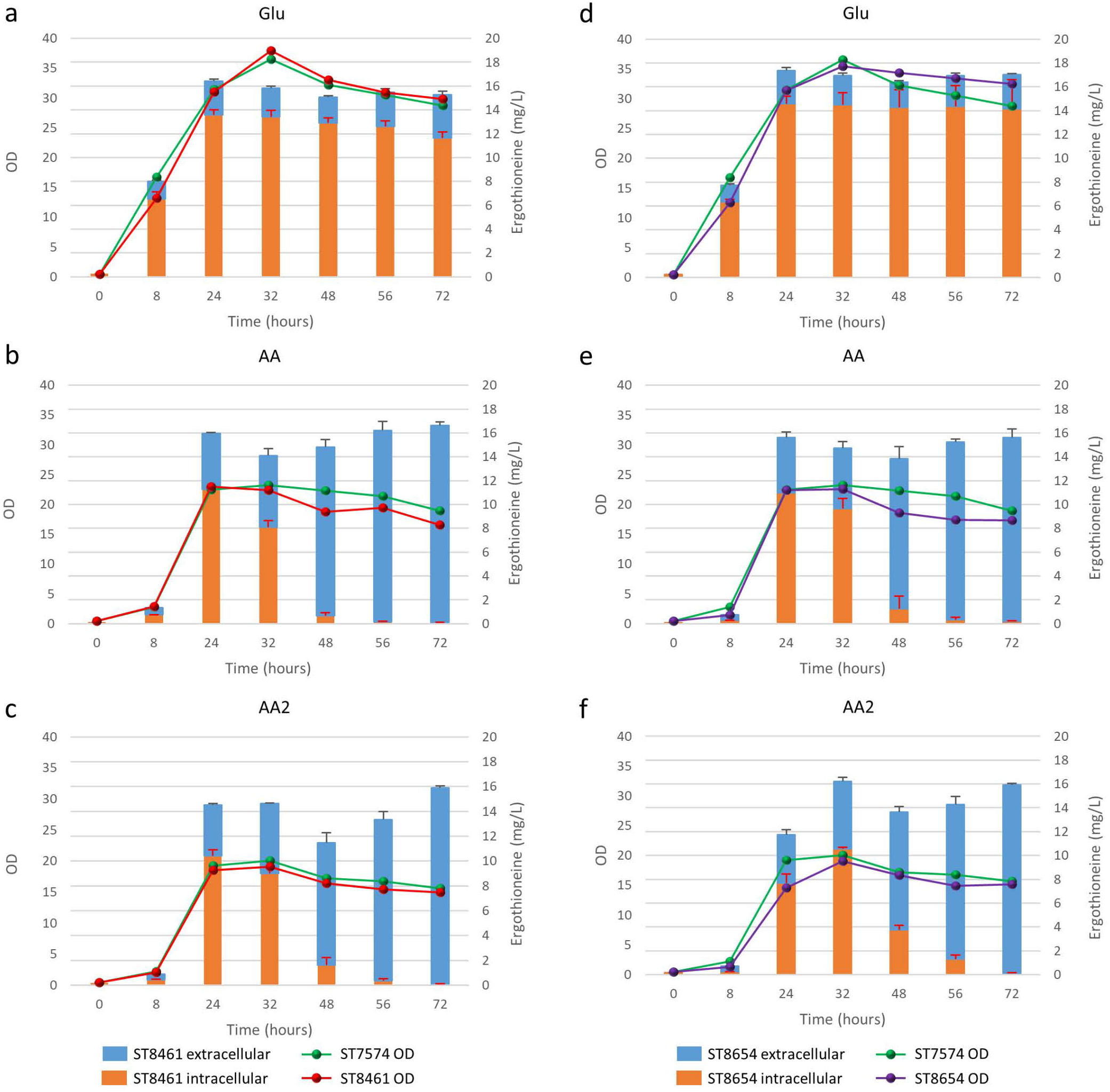
Production of ergothioneine over time in the production strain with or without transporter in different media compositions. **(A)** and **(D)** Glu: SC + 40 g/l glucose, **(B)** and **(E)** AA: SC + 40 g/l glucoe + 1 g/l His/Met/Cys, **(C)** and **(F)** AA2: SC + 40 g/l glucose + 2 g/l His/Cys/Met.

### 3.6 Medium optimization

To perform optimization of medium composition, we chose a two-level fractional factorial design, where the concentrations each of the components of the synthetic complete medium were varied 5-fold (figure 6, supplementary table 8). Strain ST8461 was cultured in SC medium (medium 65 in figure 6) to provide a baseline for the ergothioneine production during the experiment with which to compare the performance of the other media. After 48 hours, 63% of the designed media outperformed SC medium with regard to ERG titers. To be able to identify the best contributing components, the analysis was narrowed down to the eight top-performing media. Higher levels of arginine, histidine, methionine and pyridoxine were present in these media, while we could find no compound that had its concentration reduced across most or all of these media. Even though cysteine is a precursor for ergothioneine, it is not universally increased across the different media. However, as methionine can be converted into both S-adenosyl methionine and cysteine by yeast, increased levels of methionine in the medium by itself could be enough for increasing ERG titers. Pyridoxine is a precursor for pyridoxal 5’-phosphate (PLP), which binds to EgtE to facilitate the conversion of HCO to 2-(hydroxysulfanyl)hercynine in a PLP-dependent manner (Song et al., 2015). Since CpEgt2 is the fungal equivalent of EgtE, it is likely that pyridoxine has a positive effect on ERG production through CpEgt2. Interestingly, higher concentrations of arginine also increased ergothioneine production. The amino acid metabolism of *S. cerevisiae* contains many degradation and bioconversion pathways for arginine to be converted into other amino acids (Ljungdahl and Daignan-Fornier, 2012), which might contribute to the increased ERG titers.

**Figure 6:**
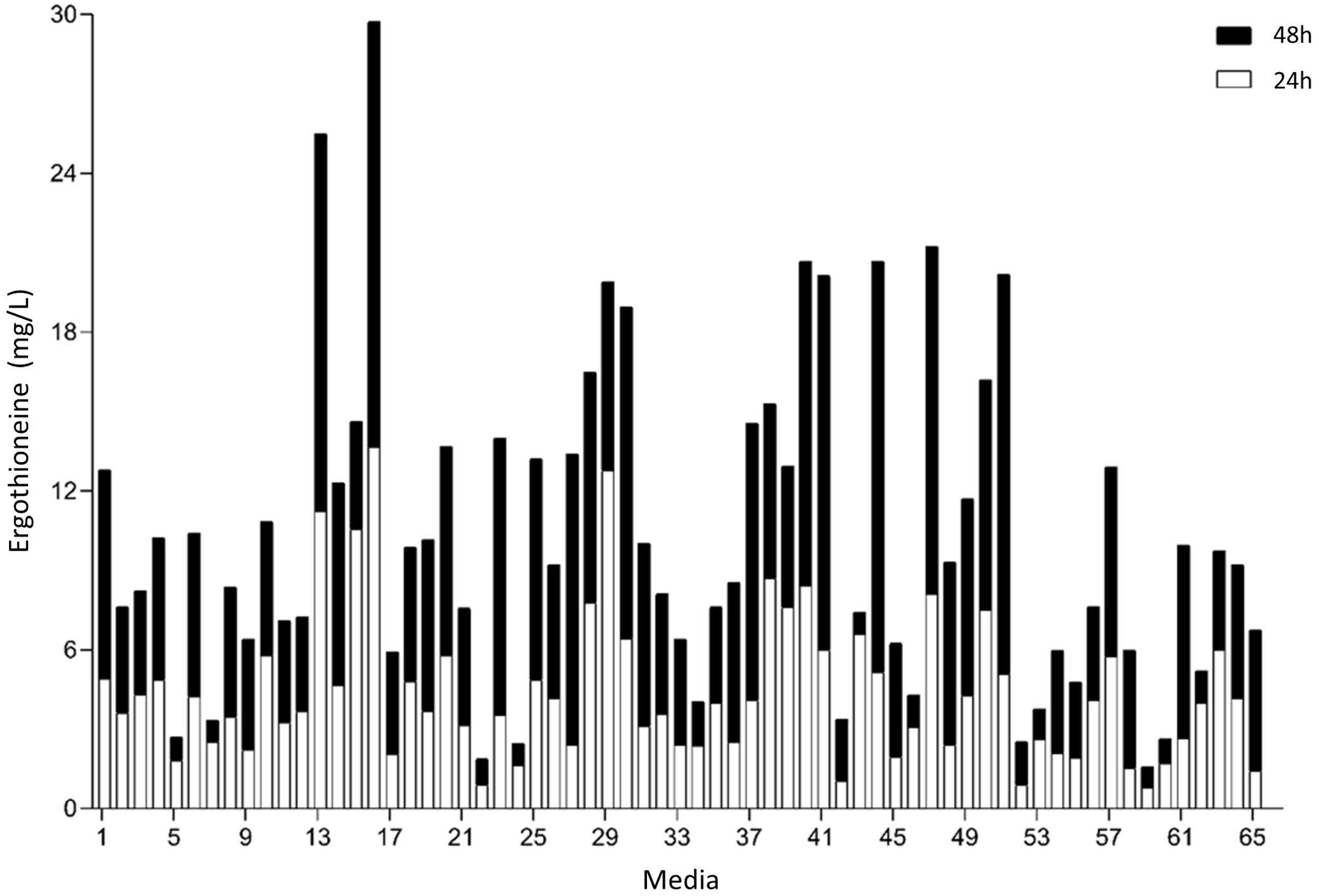
Median concentration of ergothionine produced after 24 and 48 hours. The values are additive. Open bars represent values obtained after 24 h while the closed bars represent values obtained after 48 h.

### 3.7 Enhancing the expression of ergothioneine biosynthetic genes

We recognized that the ERG-producing strain could be improved by increasing the expression of the ERG biosynthetic genes. We integrated an additional copy of NcEgt1 and/or CpEgt2 expression casseted into ST8461 (figure 7). An additional copy of NcEgt1 increased the titer by 80% and 20% for batch and simulated fed-batch medium respectively, while an additional copy of Egt2 did not increase the ergothioneine titer and even caused a decrease in simulated fed-batch medium. When additional copies of both genes were integrated, the total titer was increased by 25% on simulated fed-batch medium (ST8927). The marginal increase in titer indicates that the flux control of the pathway mainly resides elsewhere, either with the precursor supply or NcEgt1 and CpEgt2 may experience inhibition from ERG or its intermediates; this needs to be addressed through further strain engineering.

**Figure 7:**
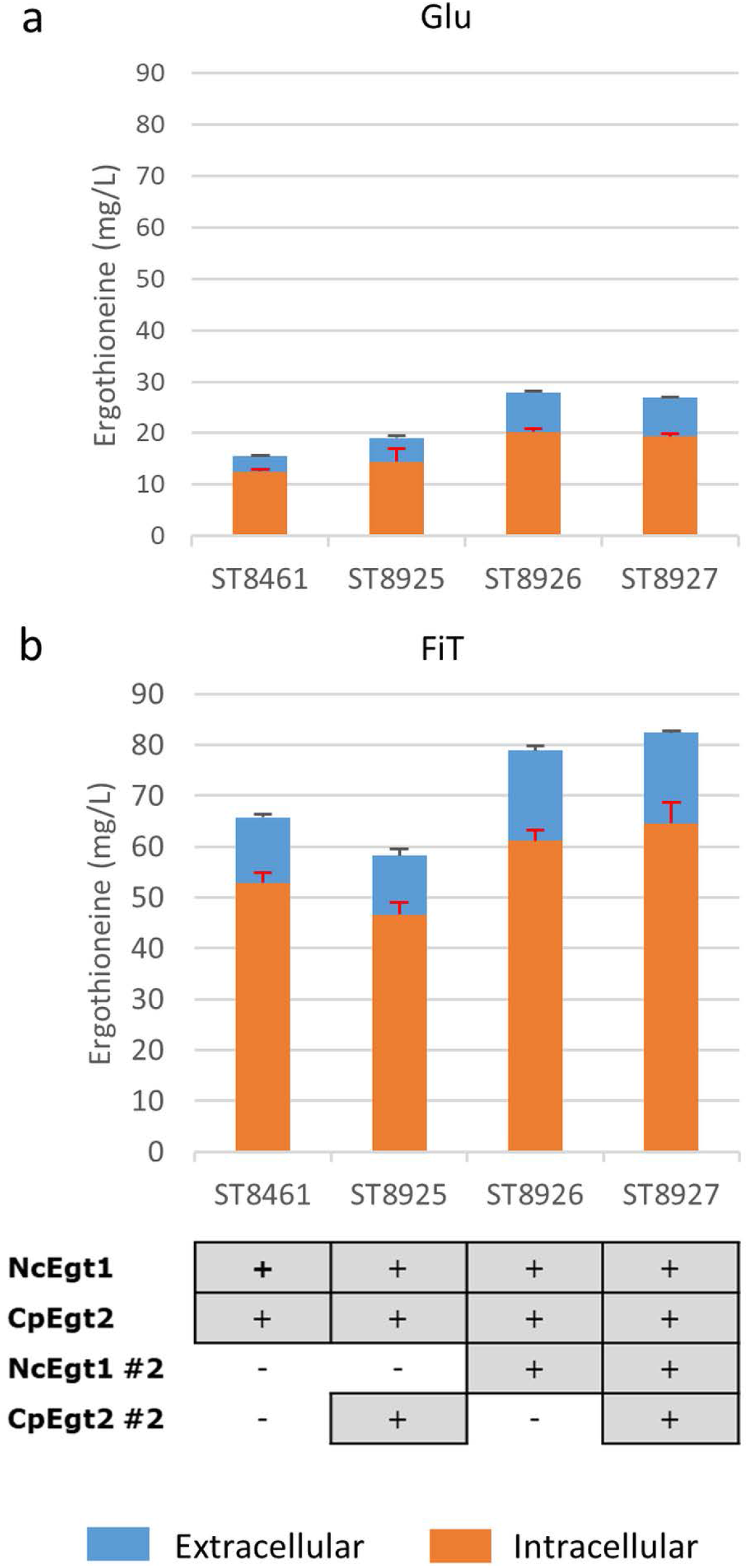
The effect of the integration of a second copy of Egt enzymes on the production of ergothioneine in the high producing strain in different media. Genomic alterations are shown as well. **(A)** Glu: SC + 40 g/l glucose **(B)** FiT: SC + 60 g/l EnPump substrate, 0.6% reagent A.

### 3.8 Ergothioneine production in controlled fed-batch fermentation

Finally, the engineered ERG-producing ST8927 strain was cultivated in bioreactors under glucose-limited fed-batch conditions. Based on the medium optimization results from section 3.6, we supplemented the fed-batch fermentation medium with arginine, histidine, methionine and pyridoxine. During the 84-hour cultivation, 598 ± 18 mg/L erg was produced, of which 59% was extracellular, from 175.0 ± 3.5 g/L glucose (figure 8). Next to that, a total of 3.2 g arginine, histidine and methionine, as well as 192 mg pyridoxine was added through the starting medium and the feeding medium. The final dry weight of biomass was 55 ± 1 g/L, and as baker’s yeast has a cell density of ~1.103 g/mL (Bryan et al., 2010), this brings the intracellular concentration of ERG to 17.7 mM, which is 11-fold higher than the extracellular concentration of 1.6 mM at the end of the fermentation. During the fermentation, at two points (after 48 hours and 72 hours), the growth of the cells started stagnating. However, when we added extra (NH_4_)_2_SO_4_, MgSO_4_, trace metal solution and vitamins, the cells began growing again. Most likely, the strain has an extra requirement for one or more of these components that was not found via the medium optimization, as the medium optimization was run under batch conditions, rather than fed-batch conditions in which yeast can reach much higher cell densities.

**Figure 8:**
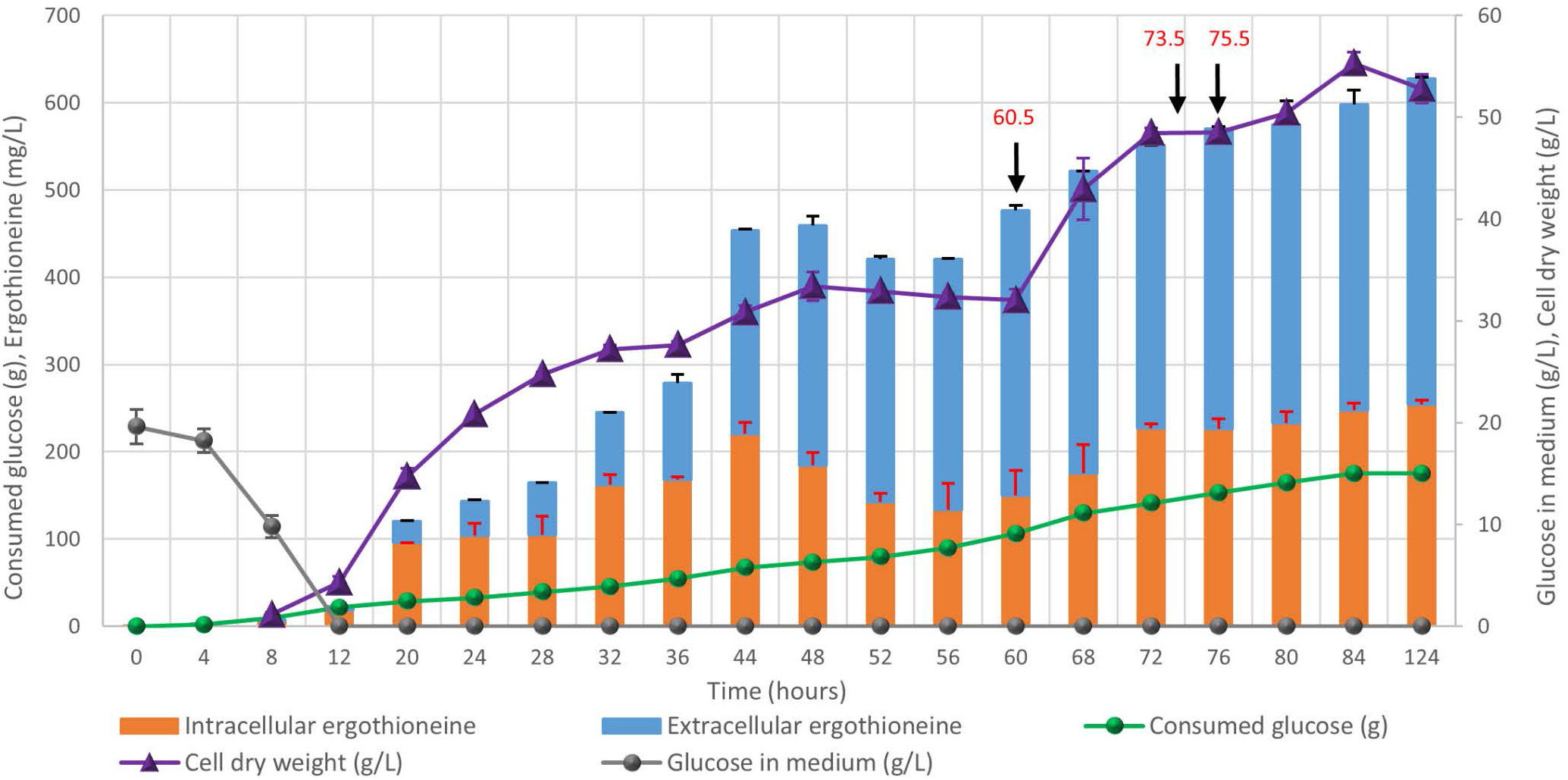
Fed-batch cultivation of ERG-producing strain ST8927. The cultivations were performed in duplicate, the average values are shown. The error bars show standard deviations. The additions of minerals, trace metals, and/or vitamins are indicated by black arrows. At 60.5 hours, we added 2 g (NH_4_)_2_SO_4_, 0.5 g MgSO_4_, 4 ml trace metals solution, and 2 ml vitamin solution, at 73.5 hours we added 0.5 g MgSO_4_, 4 ml trace metals solution 2 ml vitamin solution and at 75.5 hours we added 2 g (NH_4_)_2_SO_4_. At 60.5 and 75.5 hours, 2 g (NH_4_)_2_SO_4_ was added as a sterile 100 g/l solution. At 60.5 and 73.5 hours, 0.5 g MgSO_4_ was added as a sterile 50 g/l solution, while 4 ml sterile trace metals solution and 2 ml sterile vitamin solution were added.

## 4 Discussion

Ergothioneine is an antioxidant with many potential health benefits (Cheah and Halliwell, 2012, Ames, 2018; Halliwell et al., 2018). Furthermore, the interaction of ERG with metal ions (Hanlon, 1971) could conceivably play a role in the intracellular chaperoning of trace elements. Fermentation could provide an alternative and sustainable way for ergothioneine production compared to the current commercial processes. To this end, we have engineered *Saccharomyces cerevisiae* for the production of ergothioneine, reaching levels of 0.6 g/L. As there are many different organisms that produce ergothioneine (Genghof, 1970; Genghof et al., 1956; Genghof and Vandamme, 1964; Kalaras et al., 2017; Pfeiffer et al., 2011; Pluskal et al., 2014; Seebeck, 2010; Sheridan et al., 2016; Tanret, 1909) (supplementary table 9 and 10), we have focused on the organisms in which genes for ergothioneine production were identified. Other heterologous or from different organisms could therefore also be tested in *S. cerevisiae* to potentially find better performing combinations than those described here.

In our ERG-producing strain, as much as 59% of the ergothioneine was detected extracellularly during fermentation, even though no ERG-specific transporter has been found in yeast so far. Expression of heterologous human ergothioneine transporter or a potential ergothioneine transporter from *M. smegmatis* did not improve the secretion, however the transporters were not well expressed in the plasma membrane of *S. cerevisiae*.

While metabolic engineering of the nitrogen metabolism in yeast can lead to unexpected results, because it is so tightly regulated, engineering the strain for higher production of individual amino acids pools could lead to higher ERG titers. This is similar to the approach taken in *E. coli*, where use was made of a strain that was already overproducing cysteine (Osawa et al., 2018). In yeast, it is also possible to adopt several strategies that increase the amount of available SAM and/or methionine (Chen et al., 2016) to potentially improve ergothioneine production.

Additional to genetic manipulation, medium optimization has the benefit that it does not require extensive engineering of the strain to increase its production capabilities, while it gives a wealth of information on multiple components in the system that the strain might need for better production (Link and Weuster-Botz, 2011). However, supplementation of certain compounds can be expensive in an industrial setting and therefore the information gained can be used to lead further engineering efforts.

Ergothioneine production was only increased by a small amount following the integration of a second copy of both Egt genes. There are a number of potential explanations for this, such as a lack of precursors or (as is common (Cornishbowden et al., 1995)) possible feedback inhibition by pathway intermediates or products. Interestingly, integrating a second copy of NcEgt1 by itself, but not CpEgt2, did lead to an increase in ERG production. Indeed, in engineering *Aspergillus oryzae* (Takusagawa et al., 2019), it has been shown that hercynine accumulated following the integration of multiple copies of Egt1. This suggests that the second reaction catalyzed by the Egt1 enzyme, converting hercynine into HCO, tends to contribute more significantly to flux control.

Recently, another group managed to produce 1.3 g/L ergothioneine using *E. coli* as a production organism (Tanaka et al., 2019). They used a cysteine hyperproducing strain, which was also engineered for increased methionine production. Additionally, their ERG production genes are on a plasmid which is present in the cells with 15-20 copy numbers. During fermentation, they supplemented the medium with histidine, methionine, thiosulfate and pyridoxine, supporting the results found in our medium optimization experiment.

The fungal pathway for the biosynthesis of ERG only encompasses two enzymes compared to the five of the bacterial pathway, and eliminates the need for the use of glutamate and energy in the form of ATP. While NcEgt1 has been produced in *E. coli* for *in vitro* studies of the enzyme (Hu et al., 2014), to the best of our knowledge, the fungal pathway has to date not been used for ERG production in *E. coli*. As we have shown *S. cerevisiae* is able to produce ergothioneine using the fungal pathway, the more energetically efficient biosynthesis pathway of fungi could lead to better product yield.

In conclusion, we produced ERG with a titre of 598 ± 18 mg/L in *S. cerevisiae* expressing fungal ERG biosynthesis pathway. The study paves the way for production of ergothioneine by yeast fermentation.

## Supporting information

van der Hoek et al. Supplementary information

## Abbreviations

ERG: L-(+)-ergothioneine
HCO: S-(hercyn-2-yl)-L-cysteine S-oxide
PBS: phosphate-buffered saline
PI: propidium iodide
PLP: pyridoxal 5’-phosphate
SAM: S-adenosyl-L-methionine

## Author contributions

IB and DBK conceived the study. IB, SH, BKP, and BD designed the experiments and analyzed the data. SH, KZ performed the strain screening and transporter experiments. MB performed the nitrogen metabolism alteration experiments. SH performed the amino acid supplementation time course. BKP performed the medium optimization. The second copy integration experiments were performed by SH. The fed-batch fermentation was performed by SH, MR and JM .SH, IB and DBK wrote the manuscript. IB and DBK secured the funding and supervised the project.

## Acknowledgements

The authors would like to thank Jolanda ter Horst for her help with the fed-batch fermentation experiment.

