## Supplementary material for "Engineering the yeast *Saccharomyces cerevisiae* for the production of L-(+)-ergothioneine": van der Hoek et al. Supplementary information

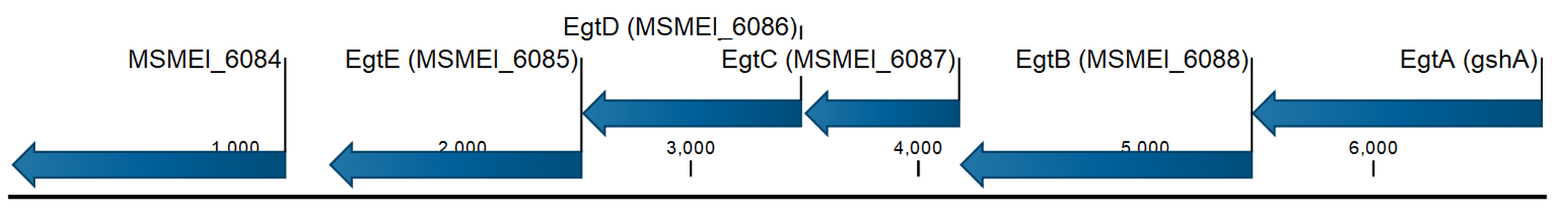

**Supplementary figure 1:** Gene cluster of ergothioneine producing genes in *Mycobacterium smegmatis*, together with MsMEI_8064, the putative ergothioneine transporter.

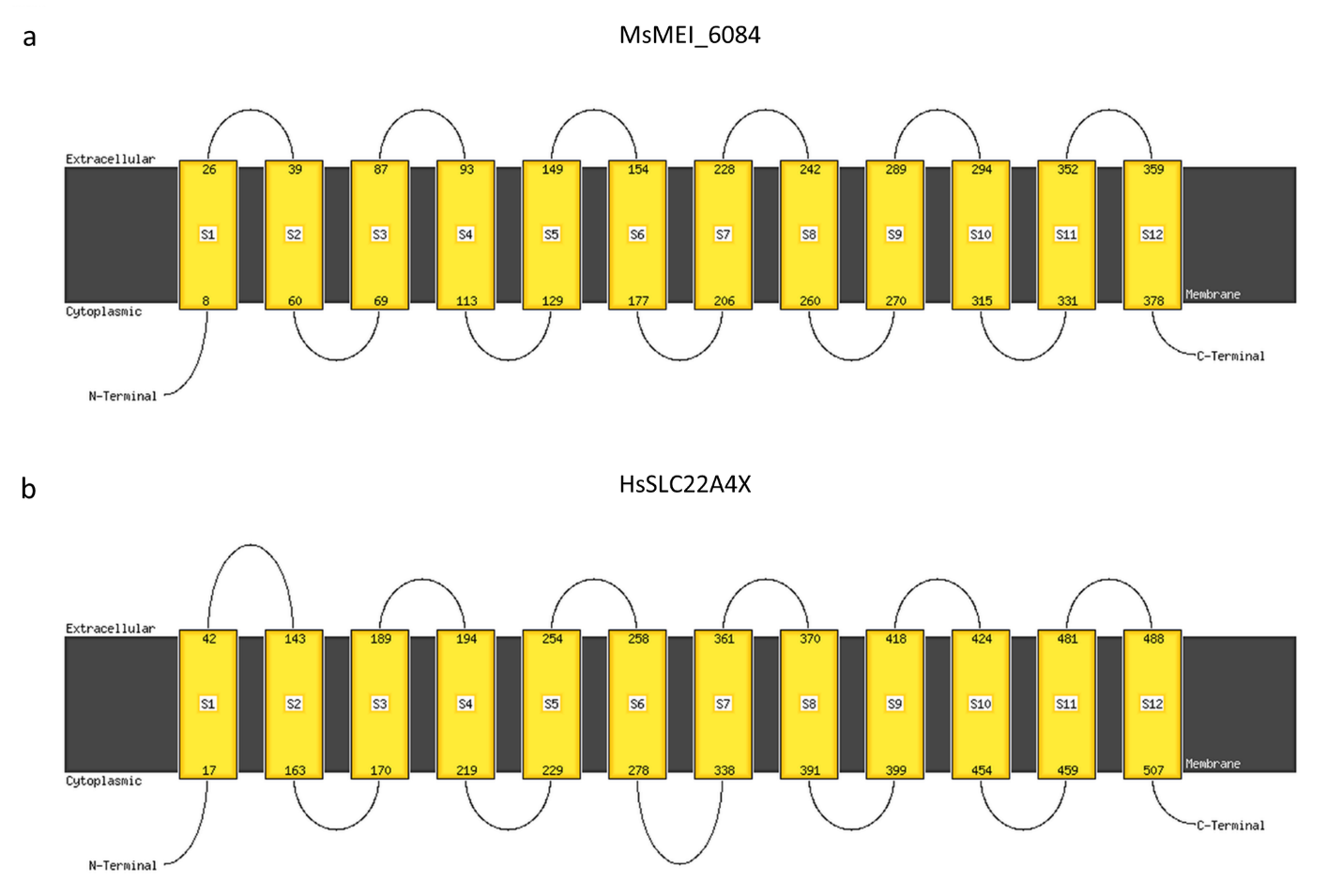

**Supplementary figure 2:** Transmembrane domain prediction by Phyre2 for **(A)** MsMEI_6084 and **(B)** HsSLC22A4X.

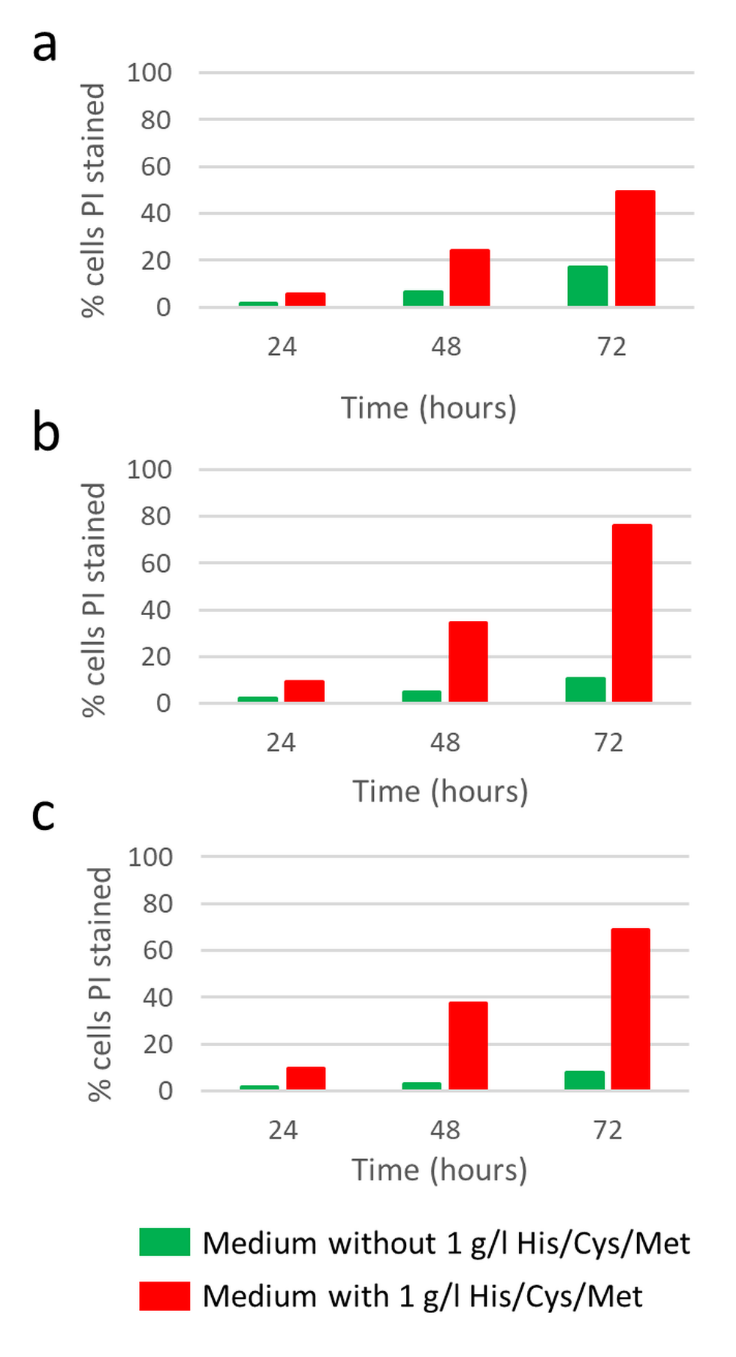

**Supplementary figure 3:** Percentage of PI stained cells for control, production strain and production strain with the transporter in media without 1 g/l histidine, cysteine and methionine versus media with 1 g/l histidine, cysteine and methionine, **(A)** Strain ST7574, **(B)** Strain ST8461, **(C)** Strain ST8654.

**Supplementary table 1:** DNA sequences and sources of the genes used in the study.

| **Protein / GenBank ID** | **DNA sequence** | **DNA source** |
| --- | --- | --- |
| MsEgtA/ AFP42520.1 | ATAATAAAAACAATGGCTTTGCCAGCTAGATCTGATTCTGGCTGTGCCGTTCCAGTCGAGTTCACTTCTGCTGAACAAGCTGCTGCCCATATTGGTGCTAACTCTTTACAAGATGGTCCAATTGGTCGTGTTGGTCTGGAAATTGAAGCTCACTGTTTCGATCTGTCTAATCCAACTCGTAGACCATCTTGGGATGAATTGTCTGCTGTCATTGCTGATGTTCCTCCATTGCCAGGAGGTTCTAGAATAACAGTGGAACCCGGAGGTGCAGTTGAATTGTCTGGTCCACCATATGATGGTCCATTGGCTGCTGTTGCTGCTTTACAAGCTGACAGGGCCGTCTTGAGGGCTGAATTTGCTAGAAGAAATTTAGGCTTGGTCTTGTTAGGTACAGATCCATTGAGACCAACGAGAAGAGTGAACCCAGGTGCTAGATATTCTGCTATGGAGCAGTTCTTCACTGCATCAGGTACTGCTGAGGCTGGTGCCGCTATGATGACTGCTACTGCATCTGTCCAAGTTAATTTGGATGCTGGTCCAAGAGATGGTTGGGCCGAGAGAGTTAGATTGGCTCATGCTTTAGGTCCCACCATGATCGCCATTACTGCTAATTCTCCAATGCTAGGTGGTCAATTTACCGGTTGGTGTTCTACAAGACAAAGAGTTTGGGGGCAATTGGATTCTGCTAGATGTGGTCCCGTTTTAGGTGTTGATGGCGACGATCCAGCCTCAGAATGGGCCAGATATGCTTTGAGAGCTCCAGTGATGTTAGTGAATTCTCCAGATGCTGTACCAGTTACTAACTGGGTCCCATTCGCTGATTGGGCTGATGGGAGAGCTGTCTTGGGTGGTAGAAGACCAACTGAAGCTGACTTGGATTATCATTTAACTACTTTATTTCCTCCAGTTAGGCCACGGAGATGGTTAGAAATTAGATATTTAGACTCGGTTCCCGACGCTTTATGGCCAGCTGCAGTTTTCACTTTAACTACTTTGTTGGATGATCCAGTTGCAGCAGAATCTGCTGCGGAAGCTACTAGACCAGTAGCTACTGCTTGGGATCGTGCTGCTAGAATGGGTTTAACTGATAGACATTTACACACCGCGGCTTTAACTTGTGTAAGATTAGCTGCTGAAAGAGCTCCGGCTGAATTGGAAGAATCTATGACATTATTAATGAGATCTGTTCAACAAAGACGGTCACCAGCTGATGATTTTTCCGATAGAGTTGTTGCTAGGGGTATCGCTGCCGCAGTTAGAGAATTGGCAAAAGGTGAATTGTGAATCTCTACTCTCTCT | Synthetic gene codon-optimized for *Saccharomyces cerevisiae* |
| MsEgtB/ WP_011731158.1 | ATGATTGCCAGAGAAACTTTGGCTGATGAATTGGCTTTGGCTAGAGAAAGAACTTTGAGATTGGTTGAATTCGATGATGCCGAATTGCATAGACAGTACAATCCATTGATGTCTCCATTGGTTTGGGACTTAGCTCATATTGGTCAACAAGAGGAATTGTGGTTGTTGAGAGATGGTAATCCAGATAGACCAGGTATGTTGGCTCCTGAAGTTGATAGATTATACGATGCCTTCGAACATTCCAGAGCTTCTAGAGTTAATTTGCCATTATTGCCACCATCTGATGCTAGAGCTTATTGTGCTACTGTTAGAGCTAAAGCTTTGGATACCTTGGATACTTTGCCAGAAGATGATCCAGGTTTTAGATTCGCCTTGGTTATCTCTCACGAAAATCAACATGACGAAACCATGTTGCAAGCCTTGAATTTGAGAGAAGGTCCACCATTATTGGATACTGGTATTCCATTGCCAGCTGGTAGACCTGGTGTTGCTGGTACTTCTGTTTTGGTTCCAGGTGGTCCATTTGTTTTGGGTGTTGATGCTTTGACTGAACCACATTCTTTGGATAACGAAAGACCAGCTCATGTTGTTGATATCCCATCTTTCAGAATTGGTAGAGTTCCAGTTACTAATGCTGAATGGCGTGAATTCATTGATGATGGTGGTTATGATCAACCTAGATGGTGGTCACCTAGAGGTTGGGCTCATAGACAAGAAGCTGGTTTGGTTGCTCCACAATTTTGGAATCCAGATGGTACTAGAACTAGATTCGGTCACATTGAAGAAATCCCAGGTGATGAACCAGTTCAACATGTTACTTTTTTCGAAGCTGAAGCTTATGCTGCTTGGGCTGGTGCTAGATTGCCAACTGAAATTGAATGGGAAAAAGCTTGTGCTTGGGATCCAGTTGCTGGTGCTCGTAGAAGATTTCCATGGGGTTCTGCTCAACCATCTGCTGCTTTAGCTAACTTAGGTGGTGATGCAAGAAGGCCAGCTCCAGTTGGTGCTTATCCAGCTGGTGCATCTGCTTATGGTGCTGAACAAATGTTGGGTGATGTTTGGGAATGGACATCTTCTCCATTAAGACCATGGCCTGGTTTTACTCCAATGATCTACGAAAGATACTCTACCCCATTCTTCGAAGGTACTACTTCTGGTGATTACAGAGTTTTGAGAGGTGGTTCATGGGCTGTTGCTCCAGGTATTTTAAGACCTTCTTTTAGAAACTGGGATCACCCAATTAGAAGGCAAATCTTTTCAGGTGTTAGATTGGCTTGGGATGTCTGA | Synthetic gene codon-optimized for *S. cerevisiae* |
| MsEgtC/ WP_011731157.1 | ATGTGCAGACATGTTGCTTGGTTGGGTGCTCCAAGATCTTTGGCTGATTTGGTTTTGGATCCACCACAAGGTTTGTTGGTTCAATCTTATGCTCCTAGAAGGCAAAAACACGGTTTGATGAATGCTGATGGTTGGGGTGCTGGTTTTTTTGATGATGAAGGTGTTGCTAGAAGATGGCGTTCTGATAAGCCTTTGTGGGGTGATGCTTCTTTTGCTTCTGTTGCTCCAGCTTTGAGATCTAGATGTGTTTTGGCTGCTGTTAGATCTGCTACTATTGGTATGCCAATTGAACCATCTGCTTCAGCTCCATTTTCTGATGGTCAATGGTTGTTGTCTCATAACGGTTTGGTTGATAGAGGTGTTTTGCCATTGACTGGTGCTGCTGAATCTACTGTTGATTCTGCTATAGTTGCTGCCTTGATTTTCTCTAGAGGTTTGGATGCTTTGGGTGCTACAATTGCTGAAGTTGGTGAATTAGATCCAAACGCCAGATTGAATATTTTGGCCGCTAATGGTTCTAGGTTGTTGGCTACTACTTGGGGTGATACTTTGTCTGTTTTACACAGACCAGATGGTGTTGTTTTAGCTTCTGAACCATATGATGATGATCCAGGTTGGTCTGATATTCCAGATAGACACTTGGTTGATGTTAGAGATGCTCATGTTGTTGTTACCCCATTGTGA | Synthetic gene codon-optimized for *S. cerevisiae* |
| MsEgtD/ WP_011731156.1 | ATGACTTTGTCCTTGGCTAATTACTTGGCTGCTGATTCTGCTGCTGAAGCTTTGAGAAGAGATGTTAGAGCTGGTTTGACTGCTGCTCCAAAATCTTTGCCACCAAAATGGTTTTATGATGCCGTTGGTTCTGATTTGTTCGATCAGATTACTAGATTGCCAGAGTACTACCCAACTAGAACTGAAGCTCAAATTTTGAGAACCAGATCCGCTGAAATTATTGCTGCTGCTGGTGCTGATACTTTGGTTGAATTAGGTTCTGGTACTTCCGAAAAGACCAGAATGTTGTTGGATGCTATGAGAGATGCCGAATTGCTGAGAAGATTCATTCCATTTGATGTTGATGCCGGTGTTTTGAGATCAGCTGGTGCAGCTATTGGTGCTGAATATCCAGGTATTGAAATTGATGCTGTTTGCGGTGATTTCGAAGAACATTTGGGTAAGATTCCACACGTTGGTAGAAGATTGGTTGTTTTCTTGGGTTCTACCATTGGTAATTTGACTCCAGCTCCAAGAGCTGAATTTTTGTCTACTTTGGCTGATACCTTGCAACCAGGTGATTCTTTGTTGTTGGGTACTGATTTGGTTAAGGATACCGGTAGATTGGTTAGAGCTTATGATGATGCAGCTGGTGTTACAGCTGCTTTTAATAGAAATGTTTTGGCCGTCGTCAACAGAGAATTGTCTGCTGATTTTGATTTGGATGCCTTCGAACATGTTGCTAAGTGGAATTCTGATGAAGAAAGGATCGAAATGTGGTTGAGAGCTAGAACTGCTCAACATGTTAGAGTTGCTGCATTGGATTTGGAAGTTGATTTTGCCGCTGGTGAAGAAATGTTGACTGAAGTTTCTTGTAAGTTCAGGCCAGAAAACGTTGTTGCTGAATTGGCTGAAGCTGGTTTAAGACAAACTCATTGGTGGACTGATCCTGCTGGTGATTTTGGTTTGTCTTTGGCTGTTAGATAA | Synthetic gene codon-optimized for *S. cerevisiae* |
| MsEgtE/ ABK70212.1 | AAAACAATGATGTTGGCTCAACAATGGAGAGATGCTAGACCAAAAGTCGCCGGTTTGCACTTAGATTCTGGTGCTTGCTCTAGACAATCTTTTGCCGTTATTGACGCAACTACTGCTCATGCTAGGCATGAAGCAGAAGTTGGTGGTTATGTTGCAGCTGAAGCCGCTACTCCAGCTTTAGATGCTGGTAGGGCTGCTGTCGCCTCTTTGATTGGTTTTGCTGCATCAGATGTTGTTTACACTTCTGGTTCTAATCACGCTATTGATTTACTATTGTCTTCTTGGCCAGGTAAAAGAACTTTAGCCTGTTTGCCCGGTGAATATGGTCCAAATTTGTCTGCTATGGCTGCAAATGGTTTTCAAGTTAGAGCTCTGCCAGTGGATGATGATGGTAGAGTTTTGGTTGATGAAGCTTCTCATGAATTGTCTGCTCATCCAGTTGCCTTAGTCCATTTGACCGCTTTGGCTTCTCATAGAGGTATTGCCCAGCCAGCAGCTGAATTGGTTGAAGCTTGTCATAACGCCGGTATCCCAGTTGTTATTGATGCTGCACAGGCATTGGGCCATTTAGATTGTAATGTTGGTGCTGATGCGGTCTATTCCTCCTCTAGAAAATGGTTGGCTGGTCCAAGGGGTGTTGGTGTACTAGCTGTTAGACCAGAATTAGCTGAAAGATTACAACCAAGAATTCCACCATCTGATTGGCCAATCCCAATGTCTGTTTTGGAAAAATTGGAATTAGGTGAGCATAACGCTGCTGCTAGAGTTGGTTTTTCTGTTGCTGTGGGTGAACATCTCGCAGCTGGACCAACTGCTGTCAGGGAAAGATTAGCTGAAGTTGGTAGATTATCTAGGCAAGTCTTGGCTGAAGTTGATGGATGGAGAGTCGTCGAACCAGTTGATCAACCAACTGCAATTACTACTTTAGAATCTACCGATGGTGCAGATCCAGCTTCTGTTAGATCTTGGTTAATCGCTGAAAGAGGTATTGTTACTACTGCTTGTGAGTTGGCTAGAGCTCCATTTGAAATGAGAACTCCAGTCCTGAGAATTTCTCCACATGTTGACGTTACAGTTGATGAATTAGAACAATTTGCTGCAGCTTTGAGAGAAGCTCCATGAAAAA | Synthetic gene codon-optimized for *S. cerevisiae* |
| NcEgt1/ XP_956324.3 | ATGCCATCTGCTGAATCTATGACTCCATCTTCTGCTTTGGGTCAATTGAAAGCTACTGGTCAACATGTCTTGTCCAAGTTGCAACAACAAACTTCCAACGCCGATATCATCGATATTAGAAGAGTTGCCGTTGAGATCAACTTGAAAACCGAAATTACCTCCATGTTCAGACCAAAAGATGGTCCAAGACAATTGCCAACCTTGTTGTTGTATAACGAAAGAGGCTTGCAGTTGTTCGAAAGAATTACTTACTTGGAAGAGTACTACTTGACCAACGACGAGATTAAGATTTTGACTAAGCACGCTACTGAAATGGCCTCTTTTATTCCATCTGGTGCCATGATTATCGAACTAGGTTCTGGTAATTTGAGGAAGGTCAACTTGTTGTTAGAAGCTTTGGATAATGCTGGTAAGGCCATTGATTATTACGCCTTGGATTTGTCCAGAGAAGAATTGGAAAGAACCTTGGCTCAAGTCCCATCTTACAAACATGTTAAGTGTCATGGTTTGTTGGGTACTTACGATGATGGTAGAGATTGGTTGAAAGCTCCAGAAAACATCAACAAGCAAAAGTGCATACTGCATCTGGGTTCTTCTATTGGTAACTTCAATAGATCTGATGCTGCCACTTTTTTGAAGGGTTTCACTGATGTTTTGGGTCCAAACGATAAGATGTTGATTGGTGTTGATGCTTGTAACGATCCAGCTAGAGTTTACCATGCTTACAATGATAAGGTTGGTATCACCCACGAATTCATCTTGAATGGTTTGAGAAACGCCAACGAAATTATTGGTGAAACCGCTTTCATTGAAGGTGATTGGAGAGTTATCGGTGAATACGTTTATGATGAAGAAGGTGGTAGACATCAAGCTTTTTATGCTCCAACTAGAGATACCATGGTTATGGGTGAATTGATCAGATCCCATGACAGAATCCAAATCGAACAGTCTCTGAAGTACTCCAAAGAGGAATCTGAAAGATTGTGGTCTACTGCTGGTTTGGAACAAGTTTCTGAATGGACTTACGGTAATGAATACGGTTTACATTTGTTGGCCAAGTCCAGAATGTCCTTCTCATTGATTCCATCAGTTTACGCTAGATCTGCTTTGCCAACTTTGGATGATTGGGAAGCTTTGTGGGCTACTTGGGATGTTGTTACTAGACAAATGTTGCCACAAGAGGAATTATTGGAGAAGCCAATCAAGTTGAGAAATGCCTGCATTTTCTACTTGGGTCATATCCCAACTTTCTTGGATATTCAGTTGACTAAGACTACCAAGCAAGCTCCATCTGAACCAGCTCATTTCTGTAAGATTTTCGAAAGGGGTATCGATCCAGATGTTGACAATCCAGAATTGTGTCATGCCCATTCTGAAATTCCAGATGAATGGCCACCAGTTGAAGAAATTTTGACTTACCAAGAAACCGTCAGATCTAGATTGAGAGGTCTATATGCTCATGGTATTGCCAACATTCCAAGAAATGTCGGTAGAGCTATTTGGGTTGGTTTCGAACATGAATTGATGCACATCGAGACTCTGTTGTACATGATGTTGCAATCTGACAAGACCTTGATTCCAACTCATATTCCAAGACCAGATTTCGATAAGTTGGCTAGAAAAGCCGAATCAGAAAGGGTTCCAAATCAATGGTTTAAGATCCCAGCTCAAGAAATCACTATTGGTTTGGATGACCCTGAAGATGGTTCCGATATTAACAAACATTACGGTTGGGATAACGAGAAGCCACCAAGAAGAGTTCAAGTTGCTGCTTTTCAAGCTCAAGGTAGACCAATTACAAACGAAGAATACGCCCAATACTTGTTGGAAAAGAACATTGATAAGTTGCCAGCTTCTTGGGCTAGATTGGATAACGAAAACATTTCTAACGGCACCACCAATTCTGTTTCTGGTCATCATTCTAACAGAACCTCCAAACAACAACTGCCATCTTCATTCTTGGAAAAAACTGCTGTTAGAACCGTTTACGGTTTGGTTCCATTGAAACATGCTTTGGATTGGCCAGTTTTTGCTTCCTATGATGAATTGGCTGGTTGTGCTGCTTATATGGGTGGTAGAATTCCAACTTTCGAAGAAACCAGATCTATCTACGCTTATGCTGATGCTCTGAAGAAGAAGAAAGAAGCTGAAAGACAACTGGGTAGAACTGTTCCAGCTGTTAATGCTCATTTGACTAACAACGGTGTTGAAATTACTCCTCCATCATCACCATCATCTGAAACTCCAGCAGAATCTTCTTCACCATCTGATTCTAACACTACCTTGATTACCACCGAGGATTTGTTCTCTGATTTGGATGGTGCTAATGTTGGTTTCCATAATTGGCATCCAATGCCTATTACTTCTAAGGGTAATACCTTGGTCGGTCAAGGTGAATTAGGTGGTGTTTGGGAATGGACATCTTCCGTTTTGAGAAAATGGGAAGGTTTTGAGCCAATGGAATTATACCCAGGTTACACTGCTGATTTCTTTGACGAAAAGCACAACATCGTTTTAGGTGGTTCATGGGCTACTCATCCAAGAATTGCTGGTAGAAAGTCTTTTGTCAACTGGTATCAAAGAAACTACCCATATGCATGGGTTGGTGCTAGAGTTGTTAGAGATTTGTGA | Synthetic gene codon-optimized for *S. cerevisiae* |
| NcEgt2/ XP_001728131.1 | ATGGTTGCTACTACTGTTGAATTGCCATTGCAACAAAAAGCTGATGCTGCTCAAACTGTTACTGGTCCATTGCCATTTGGTAACAGCTTGTTGAAAGAATTCGTTTTGGATCCAGCCTACAGAAACTTGAATCATGGTTCTTTTGGTACTATCCCATCCGCTATTCAACAGAAGTTGAGATCTTATCAAACTGCTGCTGAAGCTAGACCATGTCCATTTTTGAGATATCAAACCCCAGTTTTGTTGGACGAATCTAGAGCTGCTGTTGCTAATTTGTTGAAGGTTCCAGTTGAAACCGTTGTTTTCGTTGCTAATGCTACTATGGGTGTCAACACTGTTTTGAGAAATATCGTTTGGTCTGCTGATGGTAAGGACGAAATCTTGTACTTTGATACAATCTACGGTGCTTGCGGTAAGACCATTGATTATGTTATCGAAGATAAGAGGGGCATCGTTTCCTCTAGATGTATTCCATTGATATACCCAGCCGAAGATGATGATGTTGTTGCAGCTTTTAGAGATGCCATCAAGAAGTCTAGAGAAGAAGGTAAAAGACCAAGATTGGCCGTTATCGATGTTGTTTCTTCTATGCCAGGTGTTAGATTCCCATTCGAAGATATCGTTAAGATCTGCAAAGAGGAAGAGATCATTTCTTGCGTTGATGGTGCTCAAGGTATTGGTATGGTTGATTTGAAGATTACCGAAACCGATCCAGACTTCCTGATTTCTAATTGTCATAAGTGGTTGTTCACCCCAAGAGGTTGTGCTGTTTTTTATGTTCCAGTCAGAAACCAGCACTTGATCAGATCTACTTTGCCAACTTCTCATGGTTTCGTTCCACAAGTTGGTAATAGATTCAATCCATTGGTTCCAGCTGGTAACAAGTCTGCTTTTGTTTCTAACTTCGAATTCGTTGGTACTGTCGATAACTCTCCATTCTTCTGTGTTAAGGATGCTATTAAGTGGCGTGAAGAGGTTTTAGGTGGTGAAGAAAGAATTATGGAGTACATGACTAAGTTGGCTAGAGAAGGTGGTCAAAAGGTTGCTGAAATTTTGGGTACTAGAGTCTTGGAAAACTCTACCGGTACATTGATTAGATGCGCCATGGTTAATATTGCCTTGCCTTTTGTTGTTGGTGAAGATCCAAAAGCTCCAGTTAAGTTGACCGAAAAAGAAGAAAAAGACGTCGAAGGCTTGTACGAAATTCCACATGAAGAGGCTAATATGGCTTTCAAGTGGATGTACAACGTATTGCAAGATGAGTTCAATACCTTCGTTCCAATGACCTTTCATAGACGTAGATTTTGGGCTAGATTGTCCGCTCAAGTTTACTTGGAAATGTCTGATTTTGAATGGGCTGGCAAGACCTTAAAAGAATTGTGTGAAAGGGTTGCTAAGGGCGAGTACAAAGAATCTGCTTAA | Synthetic gene codon-optimized for *S. cerevisiae* |
| CpEgt1/ CCE33591.1 | AAAAAAAACAATGACTGCCGTTAAGCAAATTCCTGAAAGAAAGGTGTTGATAGATTCAAATCATAAGTCTCCATCAAAACCGGGTAAACATCCTAATTCTGTCATTGATATCAGGTCTAATAAGGACGATTTAAATTTACGTCATGCCCTAGTCTCATCTTTTAATCCACACGATGGAAAACCTAGGTGGCTACCTACTATGTTATTGTACGACGAAAAAGGTTTACAATTGTTTGAAGATATAACTTACTTAGATGAGTATTATTTGACTGGCTACGAAATTGAATTATTGAAGAAACATTCAGCAGAAATTGCAGCTGCTATTCCTGATGGTTCTATGGTCATCGAATTGGGCTCTGGTAATTTGAGAAAGATCTGTTTGTTGTTACAAGCCTTTGAGGATTCACATAAGTCTATCGACTACTATGCATTAGATTTATCACAAAAGGAATTAGAAAGAACTTTGAGCCATGTTCCTGACTTTAAATATGTCTCTTGTCATGGACTGCTAGGTACATATGATGATGGTGTTACATGGTTGAAACAACCAGGTATAGTCAATAAGACTAAGTGCATCATCCATCTTGGTTCGTCTATTGGGAATTTTCATAGAAATGAAGCTGCCGATTTCCTGCAGACATTTGCTGATGTAATGAAACCAGACGACTCTATGGTTATTGGTCTTGATTCATGCGGTAATCCAGAGATGTCTCGCATTCAAAGATTCATTTTGAACGGCTTATCCAATGCTAATAGCGTTTATGGCAAGGAAATATTCTATGTTCCAGATTGGAGAGTAATTGGTGAATATGTTTACGATGATGAAGGTGGCAGACACCAGGCTTTTATTTCACCTTTGAAAGAAGTCACTGCTTTAGGGTCTGTTATTAAAGCCCATGAAAGAATTAAAATTGAACAATCTTTGAAGTACTCTAAGGCCTCAGCTGACGATTTATGGAGAAATGCTGGCTTTCGAGAAACTCAAACTTGGACGAGAAACGGTGAATATGGACTACATATGTTGCAAAGAGCTGATCCGCCCTTCTCTAAGGCTCCTTCTTTGTATGCAGCTAATACTCTTCCCTCTCTTTCTGATTGGAGAGCATTGTGGTGTGCCTGGGATATTGTCACTAGAGCTATGTTGCCACAACAGGAATTGACTGAGAAACCTATAGAGTTAAGACATGCCTACATCTTTTACCTTGGTCATATTCCTACCTTCTTAGACATCCAGTTAACCAAAACATCAGCATGGGCTCCAACCTCTCCAGTTTCTTATCATGCCATTTTCGAGCGCGGCATTGATCCCGATGTTGATAACCCAGAAAAGTGTCATGATCACTCAGAGATTCCAGATGAATGGCCACCAGTCGAAGAAATTATTGCTTATCAAGATAGGGTGCGTGTTAGATTGACAGAACTGTATAAACAGGGTGTGCACACAATTACAAGAAAGGCTGCTAGAGCTATCTGGGTTTCATTTGAACATGAAGCTATGCATTTGGAAACCTTGTTGTATATGATGCTACAAAGTGATAAAGTGTTGCCACCTCCACACACTGGCGTTCCAGACTTTGAAAGAATGGCAACTAAGGCTTTCGAAGCTCGTACGCAAAATATGTGGTTCGAAATTCCAGAACAGACTATTAGTCTTGGAACAGATGATCCAGAAGATGGGGATGAAGACGTTCATTTTGGATGGGACAACGAAAAACCAGTTAGAAGAGTTAAGGTTCACGCGTTGCAAGCTCAAGGAAGACCAATTACAAATGAGGAATACGCATTATATATTTACCATACCAACTCTTCTAAACTGCCAGCATCTTGGAGTTCGTCCCCTTCATCTTCTCTGTCTAACGGCGTGTCTCATCCCAGCTCCCATAACAAGCATATTCCAACTGATTTGCCTCATTCCTTCTTGCAAGGTAAGTTTGTTAGAACCGTATATGGTTTGATACCTTTATCTTTGGCGTTGGATTGGCCTGTTCAAGCTTCTTATGATGAATTAGCTGACTGTGCATTATGGATGGGTGGAAGAATTCCAACCTTAGAGGAAGCCAGATCAATCTATGCCTTTGTTGAATCTAAAACGCAAATAGCAACAGGTAACACATTGGTCAAGAAAGTTCCTGCTGTTAATGGACACTTGGTTAATAACGGAGTTGAGGAAACTCCACCACATGAATCCTCTTCGGCAGTTGAGAATTCTTTATTCATCGACTTAGCCGGTTTGAACGTGGGTTTTAAAAGTTGGAATCCTGAACCTGTTACATCTTCTGGTACGTCTTTGGCTGGACAATCCTCTATGGGTGGTGTATGGGAGTGGACCTCTTCTGTTTTAAGACCACATGAAGGGTTCCACCCAATGGAGTTGTATCCTGGTTATACAGCCGATTTCTTTGATGAAAAACATAATATTGTTCTCGGAGGATCATGGGCTACTCATCCAAGAATAGCGGGTAGAAAAAGCTTTGTTAACTGGTATCAAAGAAACTATCCGTACGCCTGGGCTGGTGCCAGACTTGTTAAAGATGCTTGAAAAA | Synthetic gene codon-optimized for *S. cerevisiae* |
| CpEgt2/ CCE33140.1 | ATGGGTTTGTTGGAAGGTGAAGAATTGGTTTTGAGAGGTAGAGGTCAAGGTGGTGAACCTAGACCAGAAAGAGAACCAGAATTGAAGTTGGAACACGTTCCAGAAAGGGCTCCAGATGGTGAACCAGAAACTGAAGGTCAATTGGGTCCAAGAAAAGAACCTGAACATAAGTTGGAAGCTGAATCCGAACCATTGCAAGAAACTCCACAAAGAGAAGTTTTGGCTTTTGGTAGAGCTTGGAAGTCCGAATTTTTGTTTGATCCAGCTTGGAGAAACTTGAACCATGGTAGTTTTGGTACTTACCCCTTGTACATCAGAGATAAGTTGAGAGCTTATCAAGATCAAGCTGAAGCTAGACCTGATCACTTCATTAGATACGAAGAGTCCAAGTTGTTGCATAGATCTAGAGCTGCTGTTGCTAAGATAGTTAATGCTCCATTGGATACCGTTGTTTTCGTTGGTAATGCTACTGAAGGTGTCAACACTGTCTTGAGAAATTTGAGATGGGACTCCTTGGAAAAAGGTGGTCAAAAGGATGTTATCCTGTCTTTCTCTACTGTTTACGAAGCTTGTGGTAACGCTGCTGATTATATCGTTGAATACTTTGCCGGTAAGGTTGAACATAGAACCATCGAATTGGAATACCCAGTTGAAGATGCTGATGTTATTGCTGCTTTAAGAGGTGCTGCTACTCAAGTTGCTAGAGAAGGTAAAAGGGCTAGATTGGCTATGATGGATGTTGTTACTTCTAGACCAGGTGTTGTTTTTCCATGGGAAGCTGCAGTTAGAGTATGTAGAGAATTGGGTATCTTGTCCTTGGTTGATGGTGCTCAAGGTGTTGGTATGGTTAGATTGGATTTGACTGCTGCTGATCCAGATTTCTTCGTTTCTAACTGTCATAAGTGGTTGTTGGTTCCAAGAGGTTGTGCTATGTTGTATACTCCAGCTAGAACTCAATGTTTGTTGAGAACTGCTTTGGCTACTTCTCATGGTTATGTTCCACCATCTGCTGCTCCAGCTCCACCAGGTTCTAAATCTAGATATGTTGCTAACTTCGAATTCGTTGGCACTAGAGATAATGGTCCATATTTGTGTGTTGCTGATGCAATTGCTTGGAGAGAACGTGTTTGTGGTGGTGAAGAAAACATCTTGAGATACTTGTGGGCTTTGAACAAGAAGGGTATTAGAATTGTCGCTAGAGCTTTGGGTACTACCCATTTGGATAACGAAACTGAAACTTTGACCAACTGTGCTATGGGTAATGTTGCTTTGCCAATGAGAGTTGATGATGAAGATGCCTCTACTGCTTTAGATGCTGCTCCTTCTGCTGCTATTGCTGCACCAGATGTTGTTGTTGCAAGAGAAAATGTTGCATTGGTTGACAAGTGGATGAGAGAAAGATTATTCGATGACTACAAGACCTTCATGACCTTGTTCGTTATGCAAGATAGATACTGGGTTAGACTGTCTGCTCAAATCTACTTGGATGAACAAGATTATGAAGCCGCCGGTGATATTTTGAAAGCTTTGTGTGAAAGAATCAGGCGTAGAGAATATTTGGTTCCACAACCAGTTGAGTAA | Synthetic gene codon-optimized for *S. cerevisiae* |
| SpEgt1/ NP_596639.2 | ATGACAGAAATAGAAAACATTGGCGCATTAGAAGTTCTCTTCTCTCCTGAATCCATCGAGCAGAGCCTCAAACGGTGTCAACTCCCCTCCACTTTATTATACGATGAAAAAGGTTTACGACTGTTTGATGAGATTACGAATTTAAAAGAATACTACCTGTATGAAAGTGAGCTTGATATTCTGAAGAAGTTCAGCGATTCCATTGCCAACCAGTTACTGTCTCCAGATCTTCCTAACACGGTTATAGAATTAGGGTGTGGAAATATGCGCAAAACAAAACTTCTTTTAGATGCGTTTGAAAAGAAGGGCTGTGATGTGCATTTTTACGCCCTTGACCTTAATGAAGCCGAGTTGCAAAAAGGACTGCAGGAGCTTCGTCAAACTACCAATTATCAGCATGTTAAGGTGTCTGGTATTTGCGGTTGCTTTGAAAGATTGCTACAATGTTTGGACAGGTTTCGTAGTGAGCCCAATAGTCGAATTAGCATGTTGTACTTGGGTGCTTCGATTGGTAATTTTGATAGGAAATCCGCAGCATCATTTTTACGTTCGTTTGCCAGTCGTTTGAATATTCATGACAACCTTTTAATCTCCTTCGATCATAGAAACAAGGCTGAGCTAGTCCAACTAGCTTACGATGATCCTTATCGTATTACTGAAAAGTTTGAAAAGAATATTTTGGCTAGTGTCAATGCGGTTTTTGGTGAAAACCTTTTCGACGAAAATGATTGGGAATATAAAAGTGTCTACGATGAAGATCTCGGTGTTCATAGGGCCTACTTACAAGCCAAAAATGAAGTTACTGTTATTAAGGGTCCAATGTTTTTTCAATTTAAACCTAGTCATTTAATTTTGATCGAAGAAAGTTGGAAGAATAGCGATCAAGAATGTCGTCAAATCATTGAGAAAGGTGATTTTAAATTAGTCTCTAAGTATGAAAGTACGATTGCAGATTACTCGACCTATGTTATTACCAAACAATTTCCTGCTATGCTTCAACTCCCTCTTCAGCCTTGTCCTTCGTTAGCAGAATGGGATGCTCTACGCAAAGTATGGCTTTTTATTACAAATAAATTGCTTAACAAAGATAACATGTACACCGCATGGATTCCTTTGAGACATCCTCCAATTTTTTACATCGGACATGTCCCTGTTTTTAATGATATTTATCTCACAAAGATTGTCAAAAACAAAGCAACTGCTAACAAAAAACATTTTTGGGAATGGTTTCAACGTGGTATAGATCCGGACATTGAAGATCCCTCCAAGTGCCATTGGCATTCTGAAGTTCCTGAAAGCTGGCCTTCTCCTGACCAACTTCGTGAATATGAGAAAGAGTCTTGGGAATATCATATTGTAAAGTTGTGCAAAGCAATGGATGAATTGTCTACTTCTGAAAAGAGAATTCTCTGGCTTTGTTACGAACATGTAGCCATGCATGTGGAGACAACTCTTTACATCTACGTACAGTCATTTCAAAATGCAAACCAGACTGTATCAATTTGCGGATCACTTCCTGAACCAGCTGAAAAACTTACGAAAGCTCCGTTATGGGTGAATGTACCTGAAACGGAAATTGCAGTTGGTATGCCCTTGACAACACAATACACGAGTGTTGGATCAAATTTGCAATCATCCGATCTTAGTGCCCATGAAAATACAGATGAACTTTTTTATTTTGCGTGGGATAATGAGAAACCAATGAGGAAGAAACTGGTTTCTAGCTTTTCTATTGCCAATCGTCCAATTTCTAACGGTGAATATTTAGATTTTATCAATAAAAAGTCAAAAACAGAAAGGGTGTATCCAAAGCAATGGGCGGAGATTGATGGAACGCTTTACATACGAACCATGTACGGCTTATTACCCCTTGACGACTACTTGGGTTGGCCTGTTATGACTTCATACGACGATCTAAACAATTATGCGAGCTCCCAAGGATGCAGACTACCAACTGAGGATGAACTGAACTGTTTTTACGATCGGGTTCTCGAGAGAACTGATGAGCCTTATGTTAGTACCGAAGGAAAGGCAACTGGTTTTCAACAATTGCACCCTTTAGCCCTAAGTGATAATTCAAGTAATCAAATATTCACAGGAGCATGGGAATGGACAAGTACAGTTCTGGAGAAGCACGAGGATTTTGAACCTGAAGAGCTTTATCCAGATTATACACGAGATTTCTTTGATGGAAAGCATAATGTCGTTTTGGGTGGTAGCTTTGCTACGGCTACGCGCATTTCAAATAGAAGAAGCTTCAGGAACTTTTACCAAGCTGGCTATAAATATGCATGGATTGGAGCTAGACTAGTCAAAAACTAA | Genomic DNA of *Schizosaccharomyces* |
| SpEgt2/ NP_595091.1 | ATGGCTGAAAACAACGTCTACGGCCATGAAATGAAAAAGCATTTTATGCTTGATCCCGATTACGTGAATGTAAATAACGGAAGTTGTGGAACAGAATCTCTTGCTGTTTACAATAAACATGTCCAACTTTTAAAGGAAGCTCAGAGCAAGCCAGATTTTATGTGCAATGCCTATATGCCGATGTACATGGAGGCTACTCGAAATGAAGTTGCCAAGCTGATAGGCGCGGATTCAAGTAATATAGTTTTTTGCAATTCCGCTACAGATGGGATTAGTACGGTTTTGTTGACATTTCCGTGGGAACAGAATGATGAGATATTGATGCTAAATGTTGCCTATCCTACTTGTACATATGCCGCTGATTTTGCAAAGAATCAGCATAATTTACGATTAGACGTTATCGATGTTGGGGTGGAAATTGATGAAGATCTATTCCTTAAAGAAGTAGAACAGCGTTTTTTGCAGTCCAAGCCGAGAGCATTTATCTGTGATATTTTAAGTTCTATGCCCGTTATCTTGTTTCCTTGGGAAAAAGTCGTAAAGCTTTGTAAAAAGTATAATATTGTTAGCATTATTGATGGTGCTCATGCCATAGGTCATATTCCTATGAATTTGGCTAATGTTGATCCTGATTTTTTGTTTACCAATGCTCATAAATGGTTAAACTCACCAGCTGCATGCACTGTACTCTATGTCTCAGCTAAAAATCACAATCTCATCGAAGCACTTCCTCTCTCATACGGTTATGGATTAAGAGAAAAGGAATCAATTGCCGTAGATACTCTTACCAATCGGTTTGTCAATTCTTTCAAGCAAGATTTACCTAAGTTTATAGCTGTTGGTGAGGCTATTAAGTTTCGAAAATCCATTGGAGGGGAAGAAAAGATTCAACAATATTGTCATGAAATAGCTTTAAAGGGAGCCGAAATTATTTCTAAAGAACTGGGCACTTCCTTTATCAAACCTCCATACCCAGTTGCAATGGTAAACGTCGAAGTTCCCTTACGCAACATTCCCTCCATAGAAACACAGAAAGTATTTTGGCCTAAATATAATACATTCCTTCGATTTATGGAATTTAAAGGAAAATTTTACACTAGACTTAGCGGTGCGGTGTATTTAGAAGAATCAGATTTCTATTATATTGCTAAAGTTATTAAAGACTTCTGCTCTCTTTGA | Genomic DNA of *S. pombe* |
| MsMEI_6084/  AFP42515.1 | ATGCCATTTTCCCTGTACCCACTTGCAGTTGCTGTGTTCGCTATGGGAACCTCTGAATTTATGTTGGCAGGATTGGTTCCTGATATTGCAGCTGACCTCGGAGTGTCTATTGGCTCTGCTGGACTGTTGACTTCTGCTTTTGCTGTTGGAATGGTCGTGGGAGCTCCTTCTATGGCAGCTCTGACAAGAAGATGGCGCGCTAGAGTGTCTCTGAGCGCTTTTCTCCTTACCTTCGCTCTCGTGCATGTTCTGGGAGCCGTTACCACCTCTTTTGGAGTGCTGCTGGTGACAAGACTTGTGGCAGCTGTGGCTAATGCTGGATTCCTGGCCGTTGCCCTGAGTACAGCAGCAACACTGGTGCCAGCTGGAAGGCAGGGACGTGGACTTGCAGTTCTGCTTGCCGGAACAACCCTCGCAACAATTGCTGGAGTGCCAGGAGGAGCTGTGCTTGGAACAATGTTGGGATGGAGAGCAACCTTTTGGGCAATTGCTCTGCTGTGCCTGCCAGCAGTTGTTGGCATTGCAACCGCACTTCCTGCTGGCGCTGGTAGAGCCGGTTGGCCCGTTGCCGGAGCCAGTCTGTGCGATGAACTGGCCCAGCTGGGAAGAAAAAGACTGGCTCTGGCTATGCTTCTTGCTGCACTGGTCAATGCTGGAACATTTGCTACCTTTACATTTCTTGCTCCAATTGTGACAGAAAGTGCTGGACTTGGCCGCCTGTGGGTGTCTGTGGTGCTGCTTCTGTTCGGATTCGGAAGCTTTATTGGAGTGACAGTCGCCGGAAGGCTCAGCGATACAAGACCTGGAATTGTGATCGGCGCAGGTGGACCTGCTCTGCTGGCTGGATGGGCCGCTCTTGCACTTCTGTCCTCTCAGCCAGTGGCACTGTTGCCACTTGCATTCGTGCAGGGAGCACTGAGCTTTGCCGTTGGATCCACATTGATTACAAGGGTGCTGTATGAGGCATCAGCTGCCCCAACAATGGGTGGAGCTTATGCTACTGCAGCACTGAATGTCGGAGCAGCCGGTGGACCAGTCGCAGCTGCAGCCGCTCTGGGTAATCATGCAAACGTGGTGGCACCAGTTTGGGTTAGTTCCGTGATGGTGGCATTGGCTCTTCTCATCGCTGTTCCAATGCTGAAGATTGTGGCTCCAAGACCAAATCCAGCCACATCTACACCACCACTGGGAGGAAATTGCGGATAA | Synthetic gene codon-optimized for *S. cerevisiae* |
| HsSLC22A4X/  CAA71007.1 | ATGCGTGACTATGATGAGGTCATCGCATTCCTTGGCGAATGGGGACCATTCCAGCGCTTGATCTTCTTTCTGCTTAGCGCCTCTATTATCCCTAATGGCTTTAATGGTATGTCCGTTGTCTTCCTGGCCGGTACCCCTGAACATCGCTGTAGAGTGCCAGACGCCGCAAACCTGAGCAGCGCCTGGCGCAACAACTCTGTCCCTCTGAGACTGCGTGATGGCCGCGAGGTCCCCCACAGCTGTAGCCGCTACAGACTGGCCACCATCGCCAACTTCTCCGCTCTCGGACTGGAGCCAGGTCGCGATGTTGATCTGGGACAGCTGGAACAGGAGAGCTGTCTGGATGGCTGGGAGTTCAGCCAGGACGTCTACCTGTCCACCGTCGTGACCGAATGGAATCTGGTTTGTGAGGACAACTGGAAGGTGCCACTGACCACCTCCCTGTTCTTCGTTGGCGTCCTTCTGGGCTCCTTCGTGTCCGGTCAGCTGTCAGATAGATTTGGCAGGAAGAACGTTCTCTTCGCAACCATGGCTGTACAGACTGGCTTCAGCTTCCTGCAGATTTTCTCCATCAGCTGGGAGATGTTCACTGTTTTGTTTGTCATCGTGGGCATGGGCCAGATCTCCAACTATGTCGTTGCCTTCATCCTTGGAACAGAAATTCTTGGCAAGTCAGTTCGTATTATTTTCTCTACATTAGGAGTGTGCACATTTTTTGCAGTTGGCTATATGCTGCTGCCACTGTTTGCTTACTTCATCAGAGACTGGCGTATGCTGCTGCTGGCCCTGACGGTTCCTGGAGTGCTGTGTGTCCCACTGTGGTGGTTCATTCCTGAATCTCCCAGATGGCTGATCTCCCAGAGAAGATTTAGAGAGGCTGAAGATATCATCCAAAAAGCTGCAAAAATGAACAACATCGCTGTCCCAGCAGTCATTTTTGATTCTGTTGAAGAGCTGAATCCTCTGAAGCAGCAGAAAGCTTTCATTCTGGATCTGTTCAGAACTAGAAATATTGCCATTATGACCATTATGTCTTTGCTGCTTTGGATGCTGACCTCAGTGGGTTACTTTGCTCTGTCTCTGGATGCTCCTAATTTGCATGGAGATGCCTACCTGAACTGTTTCCTGTCTGCCTTGATTGAAATTCCAGCTTACATTACAGCCTGGCTGCTGTTGAGAACCCTGCCAAGGCGTTATATCATCGCTGCAGTTCTGTTCTGGGGAGGAGGTGTTCTTCTTTTCATTCAACTGGTGCCTGTCGATTATTACTTCTTGTCCATTGGTCTGGTCATGCTGGGAAAATTTGGTATCACCTCTGCTTTCTCCATGCTGTATGTCTTCACTGCTGAACTGTACCCAACCCTGGTCAGGAACATGGCTGTGGGTGTCACATCCACGGCCTCCAGAGTTGGCAGCATCATTGCCCCCTACTTTGTTTACCTCGGTGCTTACAACAGAATGCTGCCTTACATCGTCATGGGTTCTCTGACTGTCCTGATTGGAATCCTTACCCTTTTTTTCCCTGAATCCTTGGGAATGACTCTTCCAGAAACCTTAGAACAGATGCAGAAAGTGAAATGGTTCAGATCTGGAAAAAAAACAAGAGACTCAATGGAGACAGAGGAAAATCCCAAGGTTCTTATTACTGCATTCTAA | Synthetic gene codon-optimized for *S. cerevisiae* |

**Supplementary table 2:** List of primers used for cloning

| **ID** | **Name** | **Sequence 5´to 3´** |
| --- | --- | --- |
| PR-5 | pTEF1_fw | ACCTGCACUTTGTAATTAAAACTTAG |
| PR-6 | pTEF1-rv | CACGCGAUGCACACACCATAGCTTC |
| PR-1566 | <-pPGK1-pTEF1->_fw | ACCTGCACUTTGTTTTATATTTGTTG |
| PR-1567 | <-pPGK1-pTEF1->_rv | ATGACAGAUTTGTAATTAAAACTTAG |
| PR-21823 | MsEgtB_fwd | ATCTGTCAUAAAACAATGATTGCCAGAGAAACTTTGGCT |
| PR-21824 | MsEgtB_rev | CACGCGAUTCAGACATCCCAAGCCAATCTAA |
| PR-21825 | MsEgtC_fwd | ATCTGTCAUAAAACAATGTGCAGACATGTTGCTTGGTT |
| PR-21826 | MsEgtC_rev | CACGCGAUTCACAATGGGGTAACAACAACAT |
| PR-21827 | MsEgtD_fwd | AGTGCAGGUAAAACAATGACTTTGTCCTTGGCTAATTA |
| PR-21828 | MsEgtD_rev | CGTGCGAUTTATCTAACAGCCAAAGACAAACCAAAA |
| PR-21829 | CpEgt2_fwd | AGTGCAGGUAAAACAATGGGTTTGTTGGAAGGTGAAG |
| PR-21830 | CpEgt2_rev | CGTGCGAUTTACTCAACTGGTTGTGGAACCAAATATTCTCT |
| PR-21831 | NcEgt1_fwd | AGTGCAGGUAAAACAATGCCATCTGCTGAATCTATGAC |
| PR-21832 | NcEgt1_rev | CGTGCGAUTCACAAATCTCTAACAACTCTAGCACCAA |
| PR-21833 | NcEgt2_fwd | AGTGCAGGUAAAACAATGGTTGCTACTACTGTTGAATTGC |
| PR-21834 | NcEgt2_rev | CGTGCGAUTTAAGCAGATTCTTTGTACTCGCCCTTA |
| PR-21835 | SpEgt1_fwd | AGTGCAGGUAAAACAATGACAGAAATAGAAAACATTGGCGC |
| PR-21836 | SpEgt1_rev | CGTGCGAUTTAGTTTTTGACTAGTCTAGCTCCAATCCATGC |
| PR-21837 | SpEgt2_fwd | AGTGCAGGUAAAACAATGGCTGAAAACAACGTCTACG |
| PR-21838 | SpEgt2_rev | CGTGCGAUTCAAAGAGAGCAGAAGTCTTTAATAACTTTAGCAATATAATAGAAATCTGA |
| PR-22107 | MsEgtA_fwd | AGTGCAGGUAAAACAATGGCTTTGCCAGCTAGATC |
| PR-22108 | MsEgtA_rev | CGTGCGAUTCACAATTCACCTTTTGCCAATTCTCTAACTG |
| PR-22109 | MsEgtE_fwd | AGTGCAGGUAAAACAATGATGTTGGCTCAACAATGGAG |
| PR-22110 | MsEgtE_rev | CGTGCGAUTCATGGAGCTTCTCTCAAAGCTG |
| PR-22111 | CpEgt1_fwd | AGTGCAGGUAAAACAATGACTGCCGTTAAGCAAATTCC |
| PR-22112 | CpEgt1_rev | CGTGCGAUTCAAGCATCTTTAACAAGTCTGGCA |
| PR-22825 | HsSLC22A4X_MFG1_fwd | AGTGCAGGUAAAACAATGCGTGACTATGATGAGGTCATCGCATTCCTT |
| PR-22826 | HsSLC22A4X_MFG1_rev | CGTGCGAUTTAGAATGCAGTAATAAGAACCTTGGGATTTTCCTCTGTC |
| PR-22828 | MsMEI_6084_MFG1_fwd | AGTGCAGGUAAAACAATGCCATTTTCCCTGTACCCACTTGCAG |
| PR-22829 | MsMEI_6084_MFG1-rev | CGTGCGAUTTATCCGCAATTTCCTCCCAGTGGTGGT |
| PR-23559 | Tor1_gRNA | ATCCTCAGTTAACTAGCCAGGTTTTAGAGCTAGAA |
| PR-23560 | TOR1_repair_dsDNA | GTAAAGTGAAACATACATCAACCGGCTAGCAGGTTTGCATTGATTAACTGCGGTGTCATTTTTCATTTCGTGCTTTGTTTACTATTTATT |
| PR-23563 | Yih1_gRNA | AAACAGAGTCCATCACTTCCGTTTTAGAGCTAGAA |
| PR-23564 | Yih1_repair_dsDNA | TATGTAACAAGAAAAAAAAAAAAGAGAGAGGAAAGAAAAGCTCATAATCATTATGATTATGTCAAGCACCTGTAACCTTGATCACATGCG |
| PR-23861 | GFP-Nterm_fwd | AGTGCAGGUAAAACAATGTCTAAAGGTGAAGAATTATTCACTGGTGTTGTCCCA |
| PR-23862 | GFP-Nterm_rev | ATGACAGAUTTTGTACAATTCATCCATACCATGGGTAATACCAGCAG |
| PR-23863 | MsMEI_6084-Nterm_fwd | ATCTGTCAUATGCCATTTTCCCTGTACCCACTTGCA |
| PR-23864 | MsMEI_6084-Cterm_rev | ATGACAGAUTCCGCAATTTCCTCCCAGTGGTGG |
| PR-23865 | GFP-Cterm_fwd | ATCTGTCAUATGTCTAAAGGTGAAGAATTATTCACTGGTGTTGTCCCA |
| PR-23866 | GFP-Cterm_rev | CGTGCGATUTATTTGTACAATTCATCCATACCATGGGTAATACCAGCAGC |
| PR-23873 | HsSLC22A4X-Cterm_rev | ATGACAGAUTTAGAATGCAGTAATAAGAACCTTGGGATTTTCCTCTGTCTCC |
| PR-23874 | HsSLC22A4X-Nterm_fwd | ATCTGTCAUATGCGTGACTATGATGAGGTCATCGCATTCC |

**Supplementary table 3:** List of primers used for sequencing

| **ID** | **Name** | **Sequence 5´to 3´** |
| --- | --- | --- |
| PR-224 | ADH1_test_fw | GAAATTCFCTTATTTAGAAGTGTC |
| PR-225 | CYC1_test_rv | CTCCTTCCTTTTCGGTTAGAG |
| PR-21827 | MsEgtD_fwd | AGTGCAGGUAAAACAATGCCATCTGCTGAATCTATGAC |
| PR-21831 | NcEgt1_fwd | AGTGCAGGUAAAACAATGCCATCTGCTGAATCTATGAC |
| PR-21833 | NcEgt2_fwd | AGTGCAGGUAAAACAATGGTTGCTACTACTGTTGAATTGC |
| PR-22111 | CpEgt1_fwd | AGTGCAGGUAAAACAATGACTGCCGTTAAGCAAATTCC |
| PR-22277 | Seq_NcEgt1_1_fwd | TCTTTCTTCTTCTTCAGAGCA |
| PR-22278 | Seq_NcEgt1_2_fwd | TATCGAAATCTGGTCTTGGAA |
| PR-22279 | Seq_NcEgt1_3_fwd | AAAACATCAACAAGCAAAAGTGC |
| PR-22280 | Seq_NcEgt2_1_fwd | CTTAGTCATGTACTCCATAATTCTTTCTT |
| PR-22281 | Seq_SpEgt1_1_fwd | CTGTTTTTGACTTTTTATTGATAAAATCTAAATATT |
| PR-22282 | Seq_SpEgt1_2_fwd | GTTGCTTTGTTTTTGACAATCTTTGT |
| PR-22283 | Seq_SpEgt1_3_fwd | GTTTCTATGATCGAAGGAGATTAAAA |
| PR-22284 | Seq_SpEgt2_1_fwd | CCATTTATGAGCATTGGTAAACAA |
| PR-22285 | Seq_CpEgt1_1_fwd | TCTAAGGTTGGAATTCTTCCA |
| PR-22286 | Seq_CpEgt1_2_fwd | CTTCATGTTCAAATGAAACCCA |
| PR-22287 | Seq_CpEgt1_3_fwd | CTTAGAGTACTTCAAAGATTGTTCAA |
| PR-22288 | Seq_CpEgt2_1_fwd | AAACACGTTCTCTCCAAGCA |
| PR-22289 | Seq_CpEgt2_2_fwd | AACTGGGTATTCCAATTCGATG |
| PR-22290 | Seq_MsEgtB_1_rev | ACAATTCCTCTTGTTGACCAAT |
| PR-22291 | Seq_MsEgtB_2_rev | ATTGTGGAGCAACCAAACCA |
| PR-22292 | Seq_MsEgtE_1_fwd | GCTAGTACACCAACACCCCT |
| PR-22293 | Seq_MsEgtA_1_fwd | TGGAGCTCTCAAAGCATATCT |
| PR-22294 | Seq_MsEgtC_1_rev | AAAAACCAGCACCCCAACCA |
| PR-22295 | Seq_PGK1_middle_rev | AATTTCGTCACACAACAAGG |
| PR-22296 | Seq_TEF1_start_revx | TTTGAAGCTATGGTGTGTGC |
| PR-22900 | Seq_MsMEI_6084_fwd | GTAGCAAATGTTCCAGCATTG |
| PR-22901 | Seq_Hs.SLC22A4X_1_fwd | TGATATAACGCCTTGGCAGG |
| PR-22902 | Seq_Hs.SLC22A4X_2_fwd | AACGACATAGTTGGAGATCTG |
| PR-23888 | Seq_MsMEI_6084_2_fwd | GTAGCAAATGTTCCAGCATTG |
| PR-23889 | Seq_GFP_fwd | ATTAACTAAGGTATCACCTTCAAAC |
| PR-23890 | Seq_Hs.SLC22A4X_3_fwd | ACTCTACAGCGATGTTCAGG |

**Supplementary table 4:** List of BioBricks generated by PCR amplification

| **ID** | **Name** | **Primers** | **Template** |
| --- | --- | --- | --- |
| BB8 | <-pTEF1 | PR-5, PR-6 | pCfB0029 |
| BB312 | <-pPGK1-pTEF1-> | PR-1566, PR-1567 | pCfB0029 |
| BB3238 | MsEgtA | PR-22107, PR-22108 | pCfB8321 (synthetic MsEgtA gene) |
| BB3239 | MsEgtB | PR-21823, PR-21824 | pCfB8322 (synthetic MsEgtB gene) |
| BB3240 | MsEgtC | PR-21825, PR-21826 | pCfB8323 (synthetic MsEgtC gene) |
| BB3241 | MsEgtD | PR-21827, PR-21828 | pCfB8324 (synthetic MsEgtD gene) |
| BB3242 | MsEgtE | PR-22109, PR-22110 | pCfB8325 (synthetic MsEgtE gene) |
| BB3243 | NcEgt1 | PR-21831, PR-28132 | pCfB8326 (synthetic NcEgt1 gene) |
| BB3244 | NcEgt2 | PR-21833, PR-21834 | pCfB8327 (synthetic NcEgt2 gene) |
| BB3245 | CpEgt1 | PR-22111, PR-22112 | pCfB8328 (synthetic CpEgt1 gene) |
| BB3246 | CpEgt2 | PR-21829, PR-21830 | pCfB8329 (synthetic CpEgt2 gene) |
| BB3247 | SpEgt1 | PR-21835, PR-21836 | Genomic DNA from *S. pombe* |
| BB3248 | SpEgt2 | PR-21837, PR-21838 | Genomic DNA from *S. pombe* |
| BB3772 | MsMEI_6084 | PR-22828, PR-22829 | pCfB8755 (synthetic MsMEI_6084 gene) |
| BB3823 | MsMEI_6084-nterm | PR-23863, PR-22829 | pCfB8374 |
| BB3824 | MsMEI_6084-cterm | PR-22828, PR-23864 | pCfB8374 |
| BB3825 | Hs.SLC22A4X-nterm | PR-23874, PR-22826 | pCfB8375 |
| BB3826 | Hs.SLC22A4X-cterm | PR-22825, PR-23873 | pCfB8375 |
| BB3827 | yeGFP-nterm | PR-23861, PR-23862 | pCfB1914 |
| BB3828 | yeGFP-cterm | PR-23865, PR-23866 | pCfB1914 |
| BB4016 | HsSLC22A4X | PR22825, PR22826 | pCfB8388 (synthetic HsSLC22A4X gene) |

**Supplementary table 5:** List of plasmids made by USER cloning

| **ID** | **Name** | **Description** | **Template** | **Biobricks** | **Source** |
| --- | --- | --- | --- | --- | --- |
| **Basic vectors** | | | | | |
| pCfB1914 | pMeLS0025 | Plasmid carrying yeGFP for amplification of BB3827 and BB3828 |  |  | Skjoedt et al., 2016 |
| pCfB2312 | TEF1p-Cas9-CYC1t_kanMX | Episomal plasmid for Cas9 expression |  |  | Stovicek et al., 2015 |
| **Basic integrative vectors** | | | | | |
| pCfB2899 | X-2-MarkerFree | Backbone plasmid for construction of gene integration plasmids at X-2 site |  |  | Jessop-Fabre et al., 2016 |
| pCfB2903 | XI-2 | Backbone plasmid for construction of gene integration plasmids at XI-2 site |  |  | Jessop-Fabre et al., 2016 |
| pCfB3034 | X-3 | Backbone plasmid for construction of gene integration plasmids at X-3 site |  |  | Jessop-Fabre et al., 2016 |
| pCfB3039 | XII-2 | Backbone plasmid for construction of gene integration plasmids at XII-2 site |  |  | Jessop-Fabre et al., 2016 |
| **gRNA vectors** | | | | | |
| pCfB3020 | X-2 gRNA | Plasmid carrying gRNA to cut at X-2 site |  |  | Jessop-Fabre et al., 2016 |
| pCfB3042 | p-gRNA X-4 | Plasmid for gRNA at X-4 site, used as backbone to make new gRNA plasmids |  |  | Jessop-Fabre et al., 2016 |
| pCfB3043 | p-gRNA XI-1 | gRNA plasmid for targeting XI-1 site |  |  | Jessop-Fabre et al., 2016 |
| pCfB3045 | p-gRNA XI-3 | gRNA plasmid for targeting XI-3 site |  |  | Jessop-Fabre et al., 2016 |
| pCfB3051 | X-3, XI-2, XII-2 gRNA | Plasmid carrying gRNAs to cut at X-2, XI-2 and XII-2 site |  |  | Jessop-Fabre et al., 2016 |
| **Integrative vectors** | | | | | |
| pCfB8331 | pX-3-MarkerFree-NcEgt1<-TEF1 | Integration of NcEgt1 at X-3 site on *S.cerevisiae* genome | pCfB3034 | BB8, BB3243 | This study |
| pCfB8332 | pXII-2-MarkerFree-NcEgt2<-TEF1 | Integration of NcEgt2 at XII-2 site on *S.cerevisiae* genome | pCfB3039 | BB8, BB3244 | This study |
| pCfB8333 | pX-3-MarkerFree-SpEgt1<-TEF1 | Integration of SpEgt1 at X-3 site on *S.cerevisiae* genome | pCfB3034 | BB8, BB3247 | This study |
| pCfB8334 | pXII-2-MarkerFree-SpEgt2<-TEF1 | Integration of SpEgt2 at XII-2 site on *S.cerevisiae* genome | pCfB3039 | BB8, BB3248 | This study |
| pCfB8335 | pX-3-MarkerFree-CpEgt1<-TEF1 | Integration of CpEgt1 at X-3 site on *S.cerevisiae* genome | pCfB3034 | BB8, BB3245 | This study |
| pCfB8336 | pXII-2-MarkerFree-CpEgt2<-TEF1 | Integration of CpEgt2 at XII-2 site on *S.cerevisiae* genome | pCfB3039 | BB8, BB3246 | This study |
| pCfB8337 | pX-3-MarkerFree-MsEgtD<-PGK1-TEF1->MsEgtB | Integration of MsEgtD and MsEgtB at X-3 site on *S. cerevisiae* genome | pCfB3034 | BB312, BB3239, BB3241 | This study |
| pCfB8338 | pXII-2-MarkerFree-MsEgtE<-TEF1 | Integration of MsEgtE at XII-2 site on *S.cerevisiae* genome | pCfB3039 | BB8,  BB3242 | This study |
| pCfB8339 | pXI-2-MarkerFree-MsEgtA<-PGK1-TEF1->MsEgtC | Integration of MsEgtA and MsEgtC at XI-2 site on *S. cerevisiae* genome | pCfB2903 | BB312, BB3238, BB3240 | This study |
| pCfB8374 | X-2-MarkerFree-MsMEI_6084<-TEF1 | Integration of MsMEI_6084 at X-2 site on *S. cerevisiae* genome | pCfB2899 | BB8, BB3772 | This study |
| pCfB8375 | X-2-MarkerFree-Hs.SLC22A4X<-TEF1 | Integration of Hs.SLC22A4X at X-2 site on S. cerevisiae genome | pCfB2899 | BB8, BB4016 | This study |
| pCfB8800 | X-2-MarkerFree-MsMEI_6084-GFPnterm<-TEF1 | Integration of GFP-MsMEI_6084 at X-2 site on S. cerevisiae genome | pCfB2899 | BB8, BB3823, BB3827 | This study |
| pCfB8801 | X-2-MarkerFree-GFPcterm-MsMEI_6084<-TEF1 | Integration of MsMEI_6084-GFP at X-2 site on S. cerevisiae genome | pCfB2899 | BB8, BB3824, BB3828 | This study |
| pCfB8802 | X-2-MarkerFree-GFPcterm-Hs.SLC22A4X<-TEF1 | Integration of Hs.SLC22A4X-GFP at X-2 site on S. cerevisiae genome | pCfB2899 | BB8, BB3826, BB3827 | This study |
| pCfB8803 | X-2-MarkerFree-Hs.SLC22A4X-GFPnterm-TEF1 | Integration of GFP-Hs.SLC22A4X at X-2 site on S. cerevisiae genome | pCfB2899 | BB8, BB3825, BB3828 | This study |
| pCfB8804 | pXI-3-MarkerFree-NcEgt1<-pTEF1 | Integration of NcEgt1 at XI-3 site on S. cerevisiae genome | pCfB2904 | BB8, BB3243 | This study |
| pCfB8805 | pXI-1-MarkerFree-CpEgt2<-pTEF1 | Integration of CpEgt2 at XI-1 site on S.cerevisiae genome | pCfB3036 | BB8, BB3246 | This study |

**Supplementary table 6:** List of plasmids made by PCR

| **ID** | **Name** | **Description** | **Primers** | **Source** |
| --- | --- | --- | --- | --- |
| pCfB8730 | gRNA_Yih1 | gRNA plasmid targeting Yih1 for gene knock out | PR-10277,  PR-23563 | This study |
| pCfB8732 | gRNA_Tor1 | gRNA plasmid targeting TOR1 for gene knock out | PR-10277,  PR-23559 | This study |

**Supplementary table 7:** List of yeast strains

| **Strain** | **Characteristics** | **Strain specifics** | **Parent strain** | **Integration** | **Source** |
| --- | --- | --- | --- | --- | --- |
| ST1 | CEN.PK113-7D  Mata MAL2-8c SUC2 URA3 HIS3 LEU2 TRP1 | Parent strain |  |  | Peter Kötter (Goethe University, Frankfurt/Main, Germany) |
| ST7574 | CEN.PK113-7D + pCfB2312 (Cas9 plasmid) | Background strain, plasmid cured out for ERG production experiments | ST1 | pCfB2312 (no integration, episomal) | This study |
| ST8459 | NcEgt1 + NcEgt2 | Fungal pathway | ST7574 | pCfB8331, pCfB8332 | This study |
| ST8460 | NcEgt1 + SpEgt2 | Fungal pathway | ST7574 | pCfB8331, pCfB8334 | This study |
| ST8461 | NcEgt1 + CpEgt2 | Fungal pathway | ST7574 | pCfB8331, pCfB8336 | This study |
| ST8462 | SpEgt1 + SpEgt2 | Fungal pathway | ST7574 | pCfB8333, pCfB8334 | This study |
| ST8463 | SpEgt1 + NcEgt2 | Fungal pathway | ST7574 | pCfB8332, pCfB8333 | This study |
| ST8464 | SpEgt1 + CpEgt2 | Fungal pathway | ST7574 | pCfB8333, pCfB8336 | This study |
| ST8465 | CpEgt1 + CpEgt2 | Fungal pathway | ST7574 | pCfB8335, pCfB8336 | This study |
| ST8466 | CpEgt1 + NcEgt2 | Fungal pathway | ST7574 | pCfB8332, pCfB8335 | This study |
| ST8467 | CpEgt1 + SpEgt2 | Fungal pathway | ST7574 | pCfB8334, pCfB8335 | This study |
| ST8468 | MsEgtD/B + MsEgtA/C + MsEgtE | Bacterial pathway | ST7574 | pCfB8337, pCfB8338,  pCfB8339 | This study |
| ST8469 | MsEgtD/B + MsEgtA/C + NcEgt2 | Mixed pathway | ST7574 | pCfB8332, pCfB8337,  pCfB8339 | This study |
| ST8470 | MsEgtD/B + MsEgtA/C + SpEgt2 | Mixed pathway | ST7574 | pCfB8334, pCfB8337,  pCfB8339 | This study |
| ST8471 | MsEgtD/B + MsEgtA/C + CpEgt2 | Mixed pathway | ST7574 | pCfB8336, pCfB8337,  pCfB8339 | This study |
| ST8472 | NcEgt1 + MsEgtE | Mixed pathway | ST7574 | pCfB8331, pCfB8338 | This study |
| ST8473 | SpEgt1 + MsEgtE | Mixed pathway | ST7574 | pCfB8333, pCfB8338 | This study |
| ST8474 | CpEgt1 + MsEgtE | Mixed pathway | ST7574 | pCfB8335, pCfB8338 | This study |
| ST8654 | NcEgt1 + CpEgt2 + MsMEI_6084 | Fungal pathway + putative bacterial transporter from *M. smegmatis* | ST8461 | pCfB8374 | This study |
| ST8655 | NcEgt1 + CpEgt2 + HsSLC22A4X | Fungal pathway + transporter from *H. sapiens* | ST8461 | pCfB8375 | This study |
| ST8882 | NcEgt1 + CpEgt2 + MsMEI_6084 + ΔYih1 | Fungal pathway + MsMEI_6084 + Yih1 KO | ST8654 | pCfB8730,  PR-23564 | This study |
| ST8883 | NcEgt1 + CpEgt2 + MsMEI_6084 + ΔTor1 | Fungal pathway + MsMEI_6084 + Tor1 KO | ST8654 | pCfB8732,  PR-23560 | This study |
| ST8921 | NcEgt1 + CpEgt2 + GFPnterm-MsMEI_6084 | Fungal pathway + MsMEI_6084 linked to GFP at N-terminus | ST8461 | pCfB8800 | This study |
| ST8922 | NcEgt1 + CpEgt2 + MsMEI_6084-GFPcterm | Fungal pathway + MsMEI_6084 linked to GFP at C-terminus | ST8461 | pCfB8801 | This study |
| ST8923 | NcEgt1 + CpEgt2 + HsSLC22A4X- GFPcterm | Fungal pathway + HsSLC22A4X linked to GFP at C-terminus | ST8461 | pCfB8802 | This study |
| ST8924 | NcEgt1 + CpEgt2 + GFPnterm-HsSLC22A4X | Fungal pathway + HsSLC22A4X linked to GFP at N-terminus | ST8461 | pCfB8803 | This study |
| ST8925 | NcEgt1 + CpEgt2 + second copy of CpEgt2 | Fungal pathway with extra CpEgt2 | ST8461 | pCfB8805 | This study |
| ST8926 | NcEgt1 + CpEgt2 + second copy of NcEgt1 | Fungal pathway with extra NcEgt1 | ST8461 | pCfB8804 | This study |
| ST8927 | NcEgt1 + CpEgt2 + second copy of both NcEgt1 and CpEgt2 | Two copies of fungal pathway | ST8461 | pCfB8804, pCfB8805 | This study |

**Supplementary table 8:** Fractional factorial design for medium optimization. All media are based on medium #65, which is SC medium prepared as described in the Material and Methods section. We have listed the concentration of each component for medium #65.

| **Medium** | Adenine | Alanine | Ammonium sulfate | Arginine | Asparagine | Aspartate | Biotin | Boric acid | Calcium Chloride | Calcium panthothenate | Copper sulfate | Cysteine | Ferric chloride | Folic acid | Glutamate | Glutamine | Glycine | Histidine | inositol |
| --- | --- | --- | --- | --- | --- | --- | --- | --- | --- | --- | --- | --- | --- | --- | --- | --- | --- | --- | --- |
| **1** | -1 | -1 | -1 | 1 | 1 | 1 | -1 | 1 | 1 | 1 | 1 | 1 | 1 | -1 | -1 | -1 | -1 | 1 | 1 |
| **2** | 1 | -1 | -1 | 1 | -1 | 1 | 1 | -1 | 1 | 1 | -1 | -1 | 1 | -1 | 1 | -1 | -1 | 1 | -1 |
| **3** | 1 | -1 | 1 | 1 | -1 | -1 | -1 | -1 | 1 | -1 | 1 | -1 | 1 | 1 | 1 | -1 | 1 | 1 | -1 |
| **4** | 1 | 1 | 1 | 1 | 1 | 1 | 1 | 1 | 1 | 1 | 1 | 1 | 1 | 1 | 1 | 1 | 1 | 1 | 1 |
| **5** | -1 | 1 | 1 | 1 | 1 | -1 | -1 | -1 | -1 | -1 | 1 | 1 | 1 | 1 | -1 | 1 | 1 | 1 | 1 |
| **6** | 1 | -1 | -1 | 1 | -1 | -1 | 1 | -1 | 1 | 1 | 1 | 1 | -1 | 1 | 1 | -1 | -1 | 1 | -1 |
| **7** | -1 | 1 | 1 | 1 | 1 | 1 | -1 | -1 | -1 | -1 | -1 | -1 | -1 | -1 | -1 | 1 | 1 | 1 | 1 |
| **8** | -1 | -1 | -1 | -1 | 1 | -1 | -1 | -1 | 1 | -1 | -1 | 1 | -1 | -1 | -1 | -1 | -1 | -1 | 1 |
| **9** | 1 | 1 | 1 | 1 | 1 | -1 | 1 | 1 | 1 | 1 | -1 | -1 | -1 | -1 | 1 | 1 | 1 | 1 | 1 |
| **10** | 1 | -1 | 1 | 1 | -1 | 1 | -1 | -1 | 1 | -1 | -1 | 1 | -1 | -1 | 1 | -1 | 1 | 1 | -1 |
| **11** | -1 | -1 | 1 | 1 | 1 | -1 | 1 | 1 | 1 | -1 | -1 | 1 | 1 | 1 | -1 | -1 | 1 | 1 | 1 |
| **12** | 1 | -1 | -1 | 1 | 1 | 1 | 1 | -1 | -1 | -1 | -1 | -1 | -1 | 1 | 1 | -1 | -1 | 1 | 1 |
| **13** | -1 | -1 | 1 | 1 | 1 | 1 | 1 | 1 | 1 | -1 | 1 | -1 | -1 | -1 | -1 | -1 | 1 | 1 | 1 |
| **14** | 1 | -1 | 1 | -1 | -1 | 1 | -1 | 1 | 1 | 1 | -1 | -1 | -1 | 1 | 1 | -1 | 1 | -1 | -1 |
| **15** | -1 | -1 | -1 | -1 | -1 | -1 | -1 | -1 | -1 | 1 | -1 | 1 | 1 | 1 | -1 | -1 | -1 | -1 | -1 |
| **16** | -1 | -1 | 1 | 1 | -1 | 1 | 1 | 1 | -1 | 1 | 1 | -1 | 1 | 1 | -1 | -1 | 1 | 1 | -1 |
| **17** | 1 | -1 | 1 | -1 | 1 | -1 | -1 | 1 | -1 | -1 | 1 | 1 | -1 | 1 | 1 | -1 | 1 | -1 | 1 |
| **18** | -1 | 1 | 1 | 1 | -1 | 1 | -1 | -1 | 1 | 1 | -1 | -1 | 1 | 1 | -1 | 1 | 1 | 1 | -1 |
| **19** | 1 | -1 | 1 | -1 | -1 | -1 | -1 | 1 | 1 | 1 | 1 | 1 | 1 | -1 | 1 | -1 | 1 | -1 | -1 |
| **20** | 1 | 1 | -1 | -1 | -1 | 1 | -1 | -1 | -1 | -1 | 1 | 1 | 1 | 1 | 1 | 1 | -1 | -1 | -1 |
| **21** | 1 | 1 | -1 | -1 | 1 | -1 | -1 | -1 | 1 | 1 | -1 | -1 | 1 | 1 | 1 | 1 | -1 | -1 | 1 |
| **22** | 1 | 1 | 1 | -1 | -1 | 1 | 1 | -1 | -1 | 1 | 1 | -1 | -1 | 1 | 1 | 1 | 1 | -1 | -1 |
| **23** | -1 | -1 | -1 | -1 | 1 | 1 | -1 | -1 | 1 | -1 | 1 | -1 | 1 | 1 | -1 | -1 | -1 | -1 | 1 |
| **24** | -1 | 1 | 1 | -1 | 1 | 1 | -1 | 1 | -1 | 1 | -1 | 1 | -1 | 1 | -1 | 1 | 1 | -1 | 1 |
| **25** | -1 | -1 | -1 | -1 | -1 | 1 | -1 | -1 | -1 | 1 | 1 | -1 | -1 | -1 | -1 | -1 | -1 | -1 | -1 |
| **26** | 1 | -1 | -1 | -1 | -1 | -1 | 1 | 1 | 1 | -1 | 1 | -1 | -1 | -1 | 1 | -1 | -1 | -1 | -1 |
| **27** | 1 | 1 | -1 | -1 | 1 | 1 | -1 | -1 | 1 | 1 | 1 | 1 | -1 | -1 | 1 | 1 | -1 | -1 | 1 |
| **28** | -1 | 1 | 1 | -1 | -1 | -1 | -1 | 1 | 1 | -1 | 1 | -1 | -1 | 1 | -1 | 1 | 1 | -1 | -1 |
| **29** | -1 | 1 | -1 | 1 | 1 | 1 | 1 | -1 | -1 | 1 | -1 | 1 | 1 | -1 | -1 | 1 | -1 | 1 | 1 |
| **30** | 1 | 1 | -1 | 1 | 1 | 1 | -1 | 1 | 1 | -1 | 1 | -1 | -1 | 1 | 1 | 1 | -1 | 1 | 1 |
| **31** | 1 | -1 | -1 | -1 | 1 | 1 | 1 | 1 | -1 | 1 | -1 | 1 | -1 | -1 | 1 | -1 | -1 | -1 | 1 |
| **32** | -1 | -1 | 1 | 1 | -1 | -1 | 1 | 1 | -1 | 1 | -1 | 1 | -1 | -1 | -1 | -1 | 1 | 1 | -1 |
| **33** | 1 | 1 | -1 | 1 | 1 | -1 | -1 | 1 | 1 | -1 | -1 | 1 | 1 | -1 | 1 | 1 | -1 | 1 | 1 |
| **34** | 1 | -1 | -1 | -1 | -1 | 1 | 1 | 1 | 1 | -1 | -1 | 1 | 1 | 1 | 1 | -1 | -1 | -1 | -1 |
| **35** | 1 | 1 | 1 | -1 | -1 | -1 | 1 | -1 | -1 | 1 | -1 | 1 | 1 | -1 | 1 | 1 | 1 | -1 | -1 |
| **36** | -1 | -1 | 1 | -1 | 1 | 1 | 1 | -1 | 1 | 1 | 1 | 1 | -1 | 1 | -1 | -1 | 1 | -1 | 1 |
| **37** | 1 | 1 | 1 | 1 | -1 | 1 | 1 | 1 | -1 | -1 | 1 | 1 | -1 | -1 | 1 | 1 | 1 | 1 | -1 |
| **38** | -1 | -1 | 1 | -1 | -1 | 1 | 1 | -1 | -1 | -1 | 1 | 1 | 1 | -1 | -1 | -1 | 1 | -1 | -1 |
| **39** | 1 | -1 | 1 | -1 | 1 | 1 | -1 | 1 | -1 | -1 | -1 | -1 | 1 | -1 | 1 | -1 | 1 | -1 | 1 |
| **40** | 1 | -1 | 1 | 1 | 1 | 1 | -1 | -1 | -1 | 1 | -1 | 1 | 1 | 1 | 1 | -1 | 1 | 1 | 1 |
| **41** | -1 | 1 | 1 | 1 | -1 | -1 | -1 | -1 | 1 | 1 | 1 | 1 | -1 | -1 | -1 | 1 | 1 | 1 | -1 |
| **42** | -1 | -1 | 1 | -1 | -1 | -1 | 1 | -1 | -1 | -1 | -1 | -1 | -1 | 1 | -1 | -1 | 1 | -1 | -1 |
| **43** | -1 | 1 | 1 | -1 | 1 | -1 | -1 | 1 | -1 | 1 | 1 | -1 | 1 | -1 | -1 | 1 | 1 | -1 | 1 |
| **44** | 1 | -1 | 1 | 1 | 1 | -1 | -1 | -1 | -1 | 1 | 1 | -1 | -1 | -1 | 1 | -1 | 1 | 1 | 1 |
| **45** | -1 | -1 | 1 | -1 | 1 | -1 | 1 | -1 | 1 | 1 | -1 | -1 | 1 | -1 | -1 | -1 | 1 | -1 | 1 |
| **46** | -1 | 1 | -1 | -1 | 1 | 1 | 1 | 1 | -1 | -1 | -1 | -1 | 1 | 1 | -1 | 1 | -1 | -1 | 1 |
| **47** | -1 | -1 | -1 | 1 | -1 | 1 | -1 | 1 | -1 | -1 | 1 | 1 | -1 | 1 | -1 | -1 | -1 | 1 | -1 |
| **48** | 1 | -1 | -1 | -1 | 1 | -1 | 1 | 1 | -1 | 1 | 1 | -1 | 1 | 1 | 1 | -1 | -1 | -1 | 1 |
| **49** | 1 | 1 | -1 | 1 | -1 | -1 | -1 | 1 | -1 | 1 | -1 | 1 | -1 | 1 | 1 | 1 | -1 | 1 | -1 |
| **50** | 1 | 1 | 1 | -1 | 1 | 1 | 1 | -1 | 1 | -1 | 1 | -1 | 1 | -1 | 1 | 1 | 1 | -1 | 1 |
| **51** | -1 | 1 | -1 | 1 | -1 | -1 | 1 | -1 | 1 | -1 | 1 | -1 | 1 | -1 | -1 | 1 | -1 | 1 | -1 |
| **52** | 1 | 1 | 1 | 1 | -1 | -1 | 1 | 1 | -1 | -1 | -1 | -1 | 1 | 1 | 1 | 1 | 1 | 1 | -1 |
| **53** | 1 | 1 | -1 | -1 | -1 | -1 | -1 | -1 | -1 | -1 | -1 | -1 | -1 | -1 | 1 | 1 | -1 | -1 | -1 |
| **54** | -1 | -1 | -1 | 1 | -1 | -1 | -1 | 1 | -1 | -1 | -1 | -1 | 1 | -1 | -1 | -1 | -1 | 1 | -1 |
| **55** | -1 | 1 | 1 | -1 | -1 | 1 | -1 | 1 | 1 | -1 | -1 | 1 | 1 | -1 | -1 | 1 | 1 | -1 | -1 |
| **56** | 1 | 1 | -1 | 1 | -1 | 1 | -1 | 1 | -1 | 1 | 1 | -1 | 1 | -1 | 1 | 1 | -1 | 1 | -1 |
| **57** | 1 | -1 | -1 | 1 | 1 | -1 | 1 | -1 | -1 | -1 | 1 | 1 | 1 | -1 | 1 | -1 | -1 | 1 | 1 |
| **58** | -1 | 1 | -1 | -1 | -1 | -1 | 1 | 1 | 1 | 1 | 1 | 1 | 1 | 1 | -1 | 1 | -1 | -1 | -1 |
| **59** | -1 | 1 | -1 | -1 | 1 | -1 | 1 | 1 | -1 | -1 | 1 | 1 | -1 | -1 | -1 | 1 | -1 | -1 | 1 |
| **60** | -1 | 1 | -1 | 1 | 1 | -1 | 1 | -1 | -1 | 1 | 1 | -1 | -1 | 1 | -1 | 1 | -1 | 1 | 1 |
| **61** | -1 | 1 | -1 | 1 | -1 | 1 | 1 | -1 | 1 | -1 | -1 | 1 | -1 | 1 | -1 | 1 | -1 | 1 | -1 |
| **62** | -1 | 1 | -1 | -1 | -1 | 1 | 1 | 1 | 1 | 1 | -1 | -1 | -1 | -1 | -1 | 1 | -1 | -1 | -1 |
| **63** | 1 | 1 | 1 | -1 | 1 | -1 | 1 | -1 | 1 | -1 | -1 | 1 | -1 | 1 | 1 | 1 | 1 | -1 | 1 |
| **64** | -1 | -1 | -1 | 1 | 1 | -1 | -1 | 1 | 1 | 1 | -1 | -1 | -1 | 1 | -1 | -1 | -1 | 1 | 1 |
| **65** | **18 mg/L** | **76 mg/L** | **5.0 g/L** | **76 mg/L** | **76 mg/L** | **76 mg/L** | **2.0 µg/L** | **500 µg/L** | **0.1 g/L** | **400 µg/L** | **40 µg/L** | **76 mg/L** | **200 µg/L** | **2.0 µg/L** | **76 mg/L** | **76 mg/L** | **76 mg/L** | **76 mg/L** | **2.76 mg/L** |

| **Medium** | Isoleucine | Leucine | Lysine | Magnesium Sulphate | Manganese sulfate | Methionine | Nicotinic acid | p-amino benzoic acid | Phenylalanine | Potassium Iodide | Potassium Phosphate | Proline | Pyridoxine HCl | Riboflavin, | Serine | Sodium Chloride | Sodium Molybdate | Thiamine HCL |
| --- | --- | --- | --- | --- | --- | --- | --- | --- | --- | --- | --- | --- | --- | --- | --- | --- | --- | --- |
| **1** | 1 | -1 | 1 | 1 | 1 | 1 | 1 | 1 | -1 | -1 | -1 | -1 | 1 | 1 | 1 | -1 | 1 | 1 |
| **2** | 1 | 1 | -1 | 1 | 1 | -1 | -1 | 1 | -1 | 1 | -1 | -1 | 1 | -1 | 1 | 1 | -1 | 1 |
| **3** | -1 | -1 | -1 | 1 | -1 | 1 | -1 | 1 | 1 | 1 | -1 | 1 | 1 | -1 | -1 | -1 | -1 | 1 |
| **4** | 1 | 1 | 1 | 1 | 1 | 1 | 1 | 1 | 1 | 1 | 1 | 1 | 1 | 1 | 1 | 1 | 1 | 1 |
| **5** | -1 | -1 | -1 | -1 | -1 | 1 | 1 | 1 | 1 | -1 | 1 | 1 | 1 | 1 | -1 | -1 | -1 | -1 |
| **6** | -1 | 1 | -1 | 1 | 1 | 1 | 1 | -1 | 1 | 1 | -1 | -1 | 1 | -1 | -1 | 1 | -1 | 1 |
| **7** | 1 | -1 | -1 | -1 | -1 | -1 | -1 | -1 | -1 | -1 | 1 | 1 | 1 | 1 | 1 | -1 | -1 | -1 |
| **8** | -1 | -1 | -1 | 1 | -1 | -1 | 1 | -1 | -1 | -1 | -1 | -1 | -1 | 1 | -1 | -1 | -1 | 1 |
| **9** | -1 | 1 | 1 | 1 | 1 | -1 | -1 | -1 | -1 | 1 | 1 | 1 | 1 | 1 | -1 | 1 | 1 | 1 |
| **10** | 1 | -1 | -1 | 1 | -1 | -1 | 1 | -1 | -1 | 1 | -1 | 1 | 1 | -1 | 1 | -1 | -1 | 1 |
| **11** | -1 | 1 | 1 | 1 | -1 | -1 | 1 | 1 | 1 | -1 | -1 | 1 | 1 | 1 | -1 | 1 | 1 | 1 |
| **12** | 1 | 1 | -1 | -1 | -1 | -1 | -1 | -1 | 1 | 1 | -1 | -1 | 1 | 1 | 1 | 1 | -1 | -1 |
| **13** | 1 | 1 | 1 | 1 | -1 | 1 | -1 | -1 | -1 | -1 | -1 | 1 | 1 | 1 | 1 | 1 | 1 | 1 |
| **14** | 1 | -1 | 1 | 1 | 1 | -1 | -1 | -1 | 1 | 1 | -1 | 1 | -1 | -1 | 1 | -1 | 1 | 1 |
| **15** | -1 | -1 | -1 | -1 | 1 | -1 | 1 | 1 | 1 | -1 | -1 | -1 | -1 | -1 | -1 | -1 | -1 | -1 |
| **16** | 1 | 1 | 1 | -1 | 1 | 1 | -1 | 1 | 1 | -1 | -1 | 1 | 1 | -1 | 1 | 1 | 1 | -1 |
| **17** | -1 | -1 | 1 | -1 | -1 | 1 | 1 | -1 | 1 | 1 | -1 | 1 | -1 | 1 | -1 | -1 | 1 | -1 |
| **18** | 1 | -1 | -1 | 1 | 1 | -1 | -1 | 1 | 1 | -1 | 1 | 1 | 1 | -1 | 1 | -1 | -1 | 1 |
| **19** | -1 | -1 | 1 | 1 | 1 | 1 | 1 | 1 | -1 | 1 | -1 | 1 | -1 | -1 | -1 | -1 | 1 | 1 |
| **20** | 1 | -1 | -1 | -1 | -1 | 1 | 1 | 1 | 1 | 1 | 1 | -1 | -1 | -1 | 1 | -1 | -1 | -1 |
| **21** | -1 | -1 | -1 | 1 | 1 | -1 | -1 | 1 | 1 | 1 | 1 | -1 | -1 | 1 | -1 | -1 | -1 | 1 |
| **22** | 1 | 1 | -1 | -1 | 1 | 1 | -1 | -1 | 1 | 1 | 1 | 1 | -1 | -1 | 1 | 1 | -1 | -1 |
| **23** | 1 | -1 | -1 | 1 | -1 | 1 | -1 | 1 | 1 | -1 | -1 | -1 | -1 | 1 | 1 | -1 | -1 | 1 |
| **24** | 1 | -1 | 1 | -1 | 1 | -1 | 1 | -1 | 1 | -1 | 1 | 1 | -1 | 1 | 1 | -1 | 1 | -1 |
| **25** | 1 | -1 | -1 | -1 | 1 | 1 | -1 | -1 | -1 | -1 | -1 | -1 | -1 | -1 | 1 | -1 | -1 | -1 |
| **26** | -1 | 1 | 1 | 1 | -1 | 1 | -1 | -1 | -1 | 1 | -1 | -1 | -1 | -1 | -1 | 1 | 1 | 1 |
| **27** | 1 | -1 | -1 | 1 | 1 | 1 | 1 | -1 | -1 | 1 | 1 | -1 | -1 | 1 | 1 | -1 | -1 | 1 |
| **28** | -1 | -1 | 1 | 1 | -1 | 1 | -1 | -1 | 1 | -1 | 1 | 1 | -1 | -1 | -1 | -1 | 1 | 1 |
| **29** | 1 | 1 | -1 | -1 | 1 | -1 | 1 | 1 | -1 | -1 | 1 | -1 | 1 | 1 | 1 | 1 | -1 | -1 |
| **30** | 1 | -1 | 1 | 1 | -1 | 1 | -1 | -1 | 1 | 1 | 1 | -1 | 1 | 1 | 1 | -1 | 1 | 1 |
| **31** | 1 | 1 | 1 | -1 | 1 | -1 | 1 | -1 | -1 | 1 | -1 | -1 | -1 | 1 | 1 | 1 | 1 | -1 |
| **32** | -1 | 1 | 1 | -1 | 1 | -1 | 1 | -1 | -1 | -1 | -1 | 1 | 1 | -1 | -1 | 1 | 1 | -1 |
| **33** | -1 | -1 | 1 | 1 | -1 | -1 | 1 | 1 | -1 | 1 | 1 | -1 | 1 | 1 | -1 | -1 | 1 | 1 |
| **34** | 1 | 1 | 1 | 1 | -1 | -1 | 1 | 1 | 1 | 1 | -1 | -1 | -1 | -1 | 1 | 1 | 1 | 1 |
| **35** | -1 | 1 | -1 | -1 | 1 | -1 | 1 | 1 | -1 | 1 | 1 | 1 | -1 | -1 | -1 | 1 | -1 | -1 |
| **36** | 1 | 1 | -1 | 1 | 1 | 1 | 1 | -1 | 1 | -1 | -1 | 1 | -1 | 1 | 1 | 1 | -1 | 1 |
| **37** | 1 | 1 | 1 | -1 | -1 | 1 | 1 | -1 | -1 | 1 | 1 | 1 | 1 | -1 | 1 | 1 | 1 | -1 |
| **38** | 1 | 1 | -1 | -1 | -1 | 1 | 1 | 1 | -1 | -1 | -1 | 1 | -1 | -1 | 1 | 1 | -1 | -1 |
| **39** | 1 | -1 | 1 | -1 | -1 | -1 | -1 | 1 | -1 | 1 | -1 | 1 | -1 | 1 | 1 | -1 | 1 | -1 |
| **40** | 1 | -1 | -1 | -1 | 1 | -1 | 1 | 1 | 1 | 1 | -1 | 1 | 1 | 1 | 1 | -1 | -1 | -1 |
| **41** | -1 | -1 | -1 | 1 | 1 | 1 | 1 | -1 | -1 | -1 | 1 | 1 | 1 | -1 | -1 | -1 | -1 | 1 |
| **42** | -1 | 1 | -1 | -1 | -1 | -1 | -1 | -1 | 1 | -1 | -1 | 1 | -1 | -1 | -1 | 1 | -1 | -1 |
| **43** | -1 | -1 | 1 | -1 | 1 | 1 | -1 | 1 | -1 | -1 | 1 | 1 | -1 | 1 | -1 | -1 | 1 | -1 |
| **44** | -1 | -1 | -1 | -1 | 1 | 1 | -1 | -1 | -1 | 1 | -1 | 1 | 1 | 1 | -1 | -1 | -1 | -1 |
| **45** | -1 | 1 | -1 | 1 | 1 | -1 | -1 | 1 | -1 | -1 | -1 | 1 | -1 | 1 | -1 | 1 | -1 | 1 |
| **46** | 1 | 1 | 1 | -1 | -1 | -1 | -1 | 1 | 1 | -1 | 1 | -1 | -1 | 1 | 1 | 1 | 1 | -1 |
| **47** | 1 | -1 | 1 | -1 | -1 | 1 | 1 | -1 | 1 | -1 | -1 | -1 | 1 | -1 | 1 | -1 | 1 | -1 |
| **48** | -1 | 1 | 1 | -1 | 1 | 1 | -1 | 1 | 1 | 1 | -1 | -1 | -1 | 1 | -1 | 1 | 1 | -1 |
| **49** | -1 | -1 | 1 | -1 | 1 | -1 | 1 | -1 | 1 | 1 | 1 | -1 | 1 | -1 | -1 | -1 | 1 | -1 |
| **50** | 1 | 1 | -1 | 1 | -1 | 1 | -1 | 1 | -1 | 1 | 1 | 1 | -1 | 1 | 1 | 1 | -1 | 1 |
| **51** | -1 | 1 | -1 | 1 | -1 | 1 | -1 | 1 | -1 | -1 | 1 | -1 | 1 | -1 | -1 | 1 | -1 | 1 |
| **52** | -1 | 1 | 1 | -1 | -1 | -1 | -1 | 1 | 1 | 1 | 1 | 1 | 1 | -1 | -1 | 1 | 1 | -1 |
| **53** | -1 | -1 | -1 | -1 | -1 | -1 | -1 | -1 | -1 | 1 | 1 | -1 | -1 | -1 | -1 | -1 | -1 | -1 |
| **54** | -1 | -1 | 1 | -1 | -1 | -1 | -1 | 1 | -1 | -1 | -1 | -1 | 1 | -1 | -1 | -1 | 1 | -1 |
| **55** | 1 | -1 | 1 | 1 | -1 | -1 | 1 | 1 | -1 | -1 | 1 | 1 | -1 | -1 | 1 | -1 | 1 | 1 |
| **56** | 1 | -1 | 1 | -1 | 1 | 1 | -1 | 1 | -1 | 1 | 1 | -1 | 1 | -1 | 1 | -1 | 1 | -1 |
| **57** | -1 | 1 | -1 | -1 | -1 | 1 | 1 | 1 | -1 | 1 | -1 | -1 | 1 | 1 | -1 | 1 | -1 | -1 |
| **58** | -1 | 1 | 1 | 1 | 1 | 1 | 1 | 1 | 1 | -1 | 1 | -1 | -1 | -1 | -1 | 1 | 1 | 1 |
| **59** | -1 | 1 | 1 | -1 | -1 | 1 | 1 | -1 | -1 | -1 | 1 | -1 | -1 | 1 | -1 | 1 | 1 | -1 |
| **60** | -1 | 1 | -1 | -1 | 1 | 1 | -1 | -1 | 1 | -1 | 1 | -1 | 1 | 1 | -1 | 1 | -1 | -1 |
| **61** | 1 | 1 | -1 | 1 | -1 | -1 | 1 | -1 | 1 | -1 | 1 | -1 | 1 | -1 | 1 | 1 | -1 | 1 |
| **62** | 1 | 1 | 1 | 1 | 1 | -1 | -1 | -1 | -1 | -1 | 1 | -1 | -1 | -1 | 1 | 1 | 1 | 1 |
| **63** | -1 | 1 | -1 | 1 | -1 | -1 | 1 | -1 | 1 | 1 | 1 | 1 | -1 | 1 | -1 | 1 | -1 | 1 |
| **64** | -1 | -1 | 1 | 1 | 1 | -1 | -1 | -1 | 1 | -1 | -1 | -1 | 1 | 1 | -1 | -1 | 1 | 1 |
| **65** | **76 mg/L** | **380 mg/L** | **76 mg/L** | **0.5 g/L** | **400 µg/L** | **76 mg/L** | **400 µg/L** | **8.2 mg/L** | **76 mg/L** | **100 µg/L** | **1.0 g/L** | **mg/L** | **400 µg/L** | **200 µg/L** | **76 mg/L** | **0.1 g/L** | **200 µg/L** | **400 µg/L** |

| **Medium** | Threonine | Tryptophan | Tyrosine | uracil | Valine | Zinc sulfate |
| --- | --- | --- | --- | --- | --- | --- |
| **1** | 1 | 1 | 1 | 1 | -1 | -1 |
| **2** | 1 | -1 | -1 | 1 | -1 | 1 |
| **3** | -1 | 1 | -1 | 1 | 1 | 1 |
| **4** | 1 | 1 | 1 | 1 | 1 | 1 |
| **5** | -1 | 1 | 1 | 1 | 1 | -1 |
| **6** | 1 | 1 | 1 | -1 | 1 | 1 |
| **7** | -1 | -1 | -1 | -1 | -1 | -1 |
| **8** | -1 | -1 | 1 | -1 | -1 | -1 |
| **9** | 1 | -1 | -1 | -1 | -1 | 1 |
| **10** | -1 | -1 | 1 | -1 | -1 | 1 |
| **11** | -1 | -1 | 1 | 1 | 1 | -1 |
| **12** | -1 | -1 | -1 | -1 | 1 | 1 |
| **13** | -1 | 1 | -1 | -1 | -1 | -1 |
| **14** | 1 | -1 | -1 | -1 | 1 | 1 |
| **15** | 1 | -1 | 1 | 1 | 1 | -1 |
| **16** | 1 | 1 | -1 | 1 | 1 | -1 |
| **17** | -1 | 1 | 1 | -1 | 1 | 1 |
| **18** | 1 | -1 | -1 | 1 | 1 | -1 |
| **19** | 1 | 1 | 1 | 1 | -1 | 1 |
| **20** | -1 | 1 | 1 | 1 | 1 | 1 |
| **21** | 1 | -1 | -1 | 1 | 1 | 1 |
| **22** | 1 | 1 | -1 | -1 | 1 | 1 |
| **23** | -1 | 1 | -1 | 1 | 1 | -1 |
| **24** | 1 | -1 | 1 | -1 | 1 | -1 |
| **25** | 1 | 1 | -1 | -1 | -1 | -1 |
| **26** | -1 | 1 | -1 | -1 | -1 | 1 |
| **27** | 1 | 1 | 1 | -1 | -1 | 1 |
| **28** | -1 | 1 | -1 | -1 | 1 | -1 |
| **29** | 1 | -1 | 1 | 1 | -1 | -1 |
| **30** | -1 | 1 | -1 | -1 | 1 | 1 |
| **31** | 1 | -1 | 1 | -1 | -1 | 1 |
| **32** | 1 | -1 | 1 | -1 | -1 | -1 |
| **33** | -1 | -1 | 1 | 1 | -1 | 1 |
| **34** | -1 | -1 | 1 | 1 | 1 | 1 |
| **35** | 1 | -1 | 1 | 1 | -1 | 1 |
| **36** | 1 | 1 | 1 | -1 | 1 | -1 |
| **37** | -1 | 1 | 1 | -1 | -1 | 1 |
| **38** | -1 | 1 | 1 | 1 | -1 | -1 |
| **39** | -1 | -1 | -1 | 1 | -1 | 1 |
| **40** | 1 | -1 | 1 | 1 | 1 | 1 |
| **41** | 1 | 1 | 1 | -1 | -1 | -1 |
| **42** | -1 | -1 | -1 | -1 | 1 | -1 |
| **43** | 1 | 1 | -1 | 1 | -1 | -1 |
| **44** | 1 | 1 | -1 | -1 | -1 | 1 |
| **45** | 1 | -1 | -1 | 1 | -1 | -1 |
| **46** | -1 | -1 | -1 | 1 | 1 | -1 |
| **47** | -1 | 1 | 1 | -1 | 1 | -1 |
| **48** | 1 | 1 | -1 | 1 | 1 | 1 |
| **49** | 1 | -1 | 1 | -1 | 1 | 1 |
| **50** | -1 | 1 | -1 | 1 | -1 | 1 |
| **51** | -1 | 1 | -1 | 1 | -1 | -1 |
| **52** | -1 | -1 | -1 | 1 | 1 | 1 |
| **53** | -1 | -1 | -1 | -1 | -1 | 1 |
| **54** | -1 | -1 | -1 | 1 | -1 | -1 |
| **55** | -1 | -1 | 1 | 1 | -1 | -1 |
| **56** | 1 | 1 | -1 | 1 | -1 | 1 |
| **57** | -1 | 1 | 1 | 1 | -1 | 1 |
| **58** | 1 | 1 | 1 | 1 | 1 | -1 |
| **59** | -1 | 1 | 1 | -1 | -1 | -1 |
| **60** | 1 | 1 | -1 | -1 | 1 | -1 |
| **61** | -1 | -1 | 1 | -1 | 1 | -1 |
| **62** | 1 | -1 | -1 | -1 | -1 | -1 |
| **63** | -1 | -1 | 1 | -1 | 1 | 1 |
| **64** | 1 | -1 | -1 | -1 | 1 | -1 |
| **65** | **76 mg/L** | **76 mg/L** | **76 mg/L** | **76 mg/L** | **76 mg/L** | **400 µg/L** |

**Supplementary table 9:** Identity score, query length and (query cover) for genes in organisms reported to produce ergothioneine when a BLASTp search was performed against the different Egt1, Egt2 and EgtE genes. Query cover percentage in brackets behind identity score. See supplementary table 10 for Genbank accession numbers.

|  | **NcasEgt1** | **CpurEgt1** | **SpomEgt1** | **RstoEgt1** | **AnidEgt1** | **AnigEgt1** | **ProqEgt1** |
| --- | --- | --- | --- | --- | --- | --- | --- |
| **NcasEgt1** |  | 61,  842 (95) | 34,  773 (92) | 36,  876 (92) | 50,  834 (97) | 50,  835 (97) | 51,  804 (95) |
| **CpurEgt1** |  |  | 33,  773 (92) | 36,  876 (92) | 51,  834 (96) | 49,  835 (97) | 49,  804 (95) |
| **SpomEgt1** |  |  |  | 33,  876 (97) | 32,  834 (96) | 32,  835 (97) | 33,  804 (97) |
|  | **PnotEgt1** | **SsalEgt1** | **PpolEgt1** | **AoryEgt1** | **AzinEgt1** | **AcarEgt1** | **MmucEgt1** |
| **NcasEgt1** | 49,  825 (97) | 31,  1032 (87) | - | 49,  845 (97) | - | 51,  837 (96) | - |
| **CpurEgt1** | 49,  825 (96) | 33,  1032 (87) | - | 48,  845 (96) | - | 49,  837 (97) | - |
| **SpomEgt1** | 32,  825 (97) | 30,  1032 (87) | - | 32,  845 (96) | - | 32,  837 (96) | - |
|  | **NtetEgt1** | **PpulEgt1** | **PostEgt1** | **PcitEgt1** | **LedoEgt1** | **GfroEgt1** | **GlucEgt1** |
| **NcasEgt1** | 98,  876 (100) | - | 34,  853 (87) | - | 34,  555 (57) | 37,  428 (57) | - |
| **CpurEgt1** | 61,  876 (96) | - | 35,  853 (95) | - | 34,  555 (55) | 39,  428 (57) | - |
| **SpomEgt1** | 33,  876 (97) | - | 33,  853 (96) | - | 36,  555 (53) | 38,  428 (55) | - |
|  | **HeriEgt1** | **AaegEgt1** | **CcibEgt1** | **MescEgt1** |  |  |  |
| **NcasEgt1** | - | - | - | - |  |  |  |
| **CpurEgt1** | - | - | - | - |  |  |  |
| **SpomEgt1** | - | - | - | - |  |  |  |
|  | **NcasEgt2** | **CpurEgt2** | **SpomEgt2** | **RstoEgt2** | **AnidEgt2** | **AnigEgt2** | **ProqEgt2** |
| **NcasEgt2** |  | 51,  526 (94) | 31,  392 (92) | 37,  408 (93) | 43,  470 (94) | 46,  450 (94) | 44,  585 (51) |
| **CpurEgt2** |  |  | 28,  392 (83) | 33,  408 (83) | 41,  470 (83) | 43,  450 (84) | 36,  585 (45) |
| **SpomEgt2** |  |  |  | 30,  408 (96) | 30,  470 (98) | 29,  450 (97) | 28,  585 (45) |
| **Table continued on next page** | | | | | | | |
|  | **PnotEgt2** | **SsalEgt2** | **PpolEgt2** | **AoryEgt2** | **AzinEgt2** | **AcarEgt2** | **MmucEgt2** |
| **NcasEgt2** | 43,  457 (94) | 37,  678 (92) | - | 45,  461 (93) | - | 43,  433 (90) | - |
| **CpurEgt2** | 37,  457 (84) | 39,  678 (82) | - | 42,  461 (83) | - | 40,  433 (79) | - |
| **SpomEgt2** | 28,  457 (99) | 28,  678 (96) | - | 27,  461 (97) | - | 29,  433 (92) | - |
|  | **NtetEgt2** | **PpulEgt2** | **PostEgt2** | **PcitEgt2** | **LedoEgt2** | **GfroEgt2** | **GlucEgt2** |
| **NcasEgt2** | 95,  473 (100) | - | 35,  445 (94) | - | 31,  581 (92) | 31,  439 (93) | 34,  427 (89) |
| **CpurEgt2** | 51,  473 (84) | - | 32,  445 (83) | - | 30,  581 (87) | 30,  439 (83) | 32,  427 (79) |
| **SpomEgt2** | 33,  473 (97) | - | 32,  445 (97) | - | 24,  581 (98) | 29,  439 (97) | 29,  427 (94) |
|  | **HeriEgt2** | **AaegEgt2** | **CcibEgt2** | **MescEgt2** |  |  |  |
| **NcasEgt2** | - | - | 41,  255 (5) | - |  |  |  |
| **CpurEgt2** | - | - | 37,  255 (3) | - |  |  |  |
| **SpomEgt2** | - | - | 46,  255 (3) | - |  |  |  |
|  | **MsmeEgtE** | **NastEgtE** | **SalbEgtE** | **SfraEgtE** | **SgriEgtE** | **AphiEgtE** | **AfumEgtE** |
| **MsmeEgtE** |  | 34,  396 (94) | 36,  512 (46) | 36,  371 (46) | 37,  463 (46) | 34,  381 (45) | 29,  517 (46) |
|  | **MturEgtE** | **MkanEgtE** | **MintEgtE** | **MforEgtE** | **MulcEgtE** | **MbalEgtE** | **MlepEgtE** |
| **MsmeEgtE** | 66,  390 (97) | 66,  378 (98) | 62,  385 (97) | 74,  371 (100) | 64,  383 (98) | 64,  383 (98) | 39,  82 (21) |
|  | **MaviEgtE** | **MbovEgtE** | **MmarEgtE** | **MmicEgtE** | **MparEgtE** | **MphlEgtE** | **MpisEgtE** |
| **MsmeEgtE** | 61,  381 (97) | 66,  390 (97) | 64,  383 (98) | 61,  375 (97) | 61,  381 (97) | 69,  371 (99) | - |
|  | **RrhoEgtE** | **AplaEgtE** | **AmaxEgtE** | **AfloEgtE** | ***Scytonema*** | ***Oscillatoria*** | ***Rhodophyta*** |
| **MsmeEgtE** | 34,  392 (99) | 28,  388 (45) | 31,  391 (38) | 29,  389 (64) | 28-40,  396  (21-59) | 28-30,  334-390  (46-59) | 29-36,  382-437  (46-52) |

**Supplementary table 10:** Fungal and bacterial organisms reported to produce ergothioneine in literature and their Egt1/Egt2/EgtE Genbank accession numbers found through homology searches.

| **Organism (fungi)** | **Reference** | **Egt1** | **Egt2** |
| --- | --- | --- | --- |
| *Neurospora crassa (Ncas)* | Genghof et al., 1956 | XP_956324.3 | XP_001728131.1 |
| *Claviceps purpurea (Cpur)* | Tanret, 1909 | CCE33591.1 | CCE33140.1 |
| *Schizosaccharomyces pombe (Spom)* | Pluskal et al., 2014 | NP_596639.2 | NP_595091.1 |
| *Rhizopus stolonifer (Rsto)* | Genghof, 1970 | RCH97401.1 | RCI05990.1 |
| *Aspergillus nidulans (Anid)* | Genghof, 1970 | XP_680889.1 | XP_663831.1 |
| *Aspergillus niger (Anig)* | Genghof, 1970 | XP_001397117.2 | XP_001390787.2 |
| *Penicillium roqueforti (Proq)* | Genghof, 1970 | CDM31097.1 | CDM34493.1 |
| *Penicillium notatum (Pnot)* | Genghof, 1970 | KZN88090.1 | KZN85331.1 |
| *Rhodotorula glutinis (Rglu)* | Genghof, 1970 | **Not found** | **Not found** |
| *Sporobolomyces salmonicolor (Ssal)* | Genghof, 1970 | CEQ42739.1 | CEQ41088.1 |
| *Physarum polycephalum (Ppol)* | Genghof, 1970 | **Not found** | **Not found** |
| *Aspergillus oryzae (Aory)* | Genghof et al., 1956 | XP_001727309.1 | XP_001821768.1 |
| *Alternaria zinnia (Azin)* | Genghof et al., 1956 | **Not found** | **Not found** |
| *Aspergillus carbonarius (Acar)* | Genghof et al., 1956 | OOF91620.1 | OOF99450.1 |
| *Mucor mucedo (Mmuc)* | Genghof et al., 1956 | **Not found** | **Not found** |
| *Neurospora tetrasperma (Ntet)* | Genghof et al., 1956 | XP_009849693.1 | XP_009848922.1 |
| *Pullularia pullulans (Ppul)* | Genghof et al., 1956 | **Not found** | **Not found** |
| *Agaricus bisporus (Abis)* | Kalaras et al., 2017 | XP_006462499.1 | XP_006461570.1 |
| *Pleurotus ostreatus (Post)* | Kalaras et al., 2017 | KDQ26018.1 | KDQ26326.1 |
| *Pleurotus citrinopileatus (Pcit)* | Kalaras et al., 2017 | **Not found** | **Not found** |
| *Lentinula edodes (Ledo)* | Kalaras et al., 2017 | GAW05586.1 | GAV99896.1 |
| *Grifola frondosa (Gfro)* | Kalaras et al., 2017 | OBZ71212.1 | OBZ72541.1 |
| *Ganoderma lucidum (Gluc)* | Kalaras et al., 2017 | **Not found** | AUN37957.1 |
| *Hericium erinaceus (Heri)* | Kalaras et al., 2017 | **Not found** | **Not found** |
| *Agrocybe aegerita (Aaeg)* | Kalaras et al., 2017 | **Not found** | **Not found** |
| *Cantharellus cibarius (Ccib)* | Kalaras et al., 2017 | **Not found** | AWA82152.1 |
| *Boletus edulis (Bedu)* | Kalaras et al., 2017 | **Not found** | **Not found** |
| *Morchella esculenta (Mesc)* | Kalaras et al., 2017 | **Not found** | **Not found** |
| **Organism (bacteria)** | **Reference** | **EgtE** |  |
| *Mycobacterium smegmatis (Msme)* | Seebeck, 2010 | ABK70212.1 |  |
| *Nocardia asteroids (Nast)* | Genghof, 1970 | SFL89244.1 |  |
| *Streptomyces albus (Salb)* | Genghof, 1970 | WP_030543061.1 |  |
| *Streptomyces fradiae (Sfra)* | Genghof, 1970 | WP_070159474.1 |  |
| *Streptomyces griseus (Sgri)* | Genghof, 1970 | WP_030852754.1 |  |
| *Actinoplanes philippinensis (Aphi)* | Genghof, 1970 | WP_093610803.1 |  |
| *Aspergillus fumigatus (Afum)* | Sheridan et al., 2016 | XP_754202.1 |  |
| *Mycobacterium tuberculosis (Mtur)* | Genghof and Vandamme, 1964 | WP_079029600.1 |  |
| *Mycobacterium kansasii (Mkan)* | Genghof and Vandamme, 1964 | WP_103802346.1 |  |
| *Mycobacterium intracellulare (Mint)* | Genghof and Vandamme, 1964 | WP_014941167.1 |  |
| *Mycobacterium forfuitum (Mfor)* | Genghof and Vandamme, 1964 | WP_076203140.1 |  |
| *Mycobacterium ulcerans (Mulc)* | Genghof and Vandamme, 1964 | WP_096369529.1 |  |
| *Mycobacterium balnei (Mbal)* | Genghof and Vandamme, 1964 | WP_117431391.1 |  |
| *Mycobacterium leprae (Mlep)* | Genghof and Vandamme, 1964 | WP_041323321.1 |  |
| *Mycobacterium avium (Mavi)* | Genghof and Vandamme, 1964 | WP_044543419.1 |  |
| *Mycobacterium bovis (Mbov)* | Genghof and Vandamme, 1964 | YP_009361087.1 |  |
| *Mycobacterium marinum (Mmar)* | Genghof and Vandamme, 1964 | WP_117431391.1 |  |
| *Mycobacterium microti (Mmic)* | Genghof and Vandamme, 1964 | PLV46245.1 |  |
| *Mycobacterium paratuberculosis (Mpar)* | Genghof and Vandamme, 1964 | AAS02619.1 |  |
| *Mycobacterium phlei (Mphl)* | Genghof and Vandamme, 1964 | WP_003888643.1 |  |
| *Mycobacterium piscinum (Mpis)* | Genghof and Vandamme, 1964 | **Not found** |  |
| *Rhodococcus rhodocrous (Rrho)*  ***Reclassified Mycobacterium rhodocrous*** | Genghof and Vandamme, 1964 | WP_006938916.1  *Multispecies* |  |
| *Arthrospira platensis (Apla)* | Pfeiffer et al., 2011 | WP_062945872.1 |  |
| *Arthrospira maxima (Amax)* | Pfeiffer et al., 2011 | WP_006621917.1  *Multispecies* |  |
| *Aphanizomenon flos-aquae (Aflo)* | Pfeiffer et al., 2011 | OBQ29810.1 |  |
| *Scytonema* sp.  **Genus wide search based on reference** | Pfeiffer et al., 2011 | WP_073633333.1  WP_096565387.1 |  |
| *Oscillatoria* sp.  **Genus wide search based on reference** | Pfeiffer et al., 2011 | WP_044196545.1  WP_015175683.1 |  |
| *Rhodophyta* sp.  **Genus wide search based on reference** | Pfeiffer et al., 2011 | OSX68822.1  PXF47457.1 |  |
